# Molecular characterization of the effector-immunity pair Rhs2-SciX from *Salmonella* Typhimurium

**DOI:** 10.64898/2026.05.10.723254

**Authors:** Neil Lorente Cobo, Shubham Goyal, David Davidson, Gerd Prehna

## Abstract

The type VI secretion system (T6SS) is a dynamic nanomachine used by bacteria to compete for space and nutrients. To kill rival bacteria, the T6SS secretes toxic effector proteins directly into adjacent cells in a contact-dependent manner. Effectors have diverse biochemical functions that arise from subtle structural modifications to related enzyme folds. To protect from self-intoxication, effectors are encoded with a cognate immunity protein in effector-immunity pairs. Immunity proteins inhibit toxicity by directly binding the effector active site and are as structurally diverse as their effectors. *Salmonella* Typhimurium has several known effectors of diverse biochemical functions, however the effector-immunity pair Rhs2-SciX remains largely unstudied. Here, we take a structural approach and show that Rhs2 is a BECR family nuclease through modeling and toxicity assays. Furthermore, we solve an X-ray crystal structure of SciX and study the solution-state properties of the immunity protein to gain insight into the binding and inhibition mechanism of Rhs2. Finally, we discover that Rhs2 binds directly to EF-Tu suggesting that Rhs2 inhibits translation to cause cell death. Our work probes the structure and function of the effector-immunity pair Rhs2-SciX, demonstrates the biochemical diversity of *Salmonella* T6SS effectors, and highlights the conserved T6SS toxicity strategy of binding EF-Tu.

## INTRODUCTION

Bacteria compete over the availability of both nutrients and space. For bacterial pathogens, they must also respond to challenges from the host^1–4^. Due to these pressures, bacteria have evolved numerous competition strategies that include diverse protein secretion systems. Bacterial secretion systems are dynamic nanomachines designed for the translocation of proteins into the environment, directly into rival bacterial cells, or even into the cells of a eukaryotic host^3,5,6^. For Gram-negative bacteria, the type VI secretion system (T6SS) is a primary mechanism of bacterial competition. The T6SS forcibly injects toxic effectors into adjacent cells in a contact-dependent manner to kill rival bacteria^7–9^. This function ultimately provides T6SS harboring bacteria a competitive advantage^9,10^.

The T6SS is structurally related to the T4 bacteriophage and consists of a needle surrounded by a contractile sheath, a baseplate, and membrane complex that spans both the inner and outer membranes^11–13^. The needle is assembled from the polymerization of hexameric <u>H</u>emolysin <u>c</u>o-regulated <u>p</u>roteins (Hcp) which are surrounded by a contractile sheath that provides energy for secretion^13,14^. Hcp needle assembly is nucleated when a <u>v</u>aline <u>g</u>lycine repeat protein G (VgrG) is properly loaded into the baseplate environment on the bacterial inner membrane^15–17^. VgrG forms the “spike” tip of the Hcp needle^18,19^, and also functions a as modular platform for effector loading^20–22^. Additionally, VgrG proteins can be sharpened by proline-alanine-alanine-arginine (PAAR) domain-containing domains^17,23,24^ and PAAR-like proteins such as PIPY-domains^25^. PAAR and PAAR-like domains serve as adaptors for effectors that bind VgrGs, and regulate the loading of VgrG-effector complexes into the T6SS^17^. Finally, the membrane spanning complex positions the T6SS needle for ejection and undergoes a conformational change as it provides the path for Hcp-VgrG-effector secretion^26,27^.

T6SS effectors have highly diverse biochemical functions and are known to kill both prokaryotic and eukaryotic cells^10,28,29^. Effectors have been shown to structurally modify or hydrolyze the peptidoglycan^30^, inhibit different steps in translation^31,32^, alter pools of essential metabolites such as NAD+/NADH^33^, cross-link actin^34^, and form pores to depolarize membranes^35^. For anti-bacterial effectors, each effector is typically co-expressed with a cognate immunity protein to prevent self-killing^10^. Immunity proteins are equally as diverse as their effectors^10,36^, and commonly inhibit effector function by binding directly to the active site of the effector^31,37^.

A prime example of effector diversity is *Salmonella*, where studies found that *Salmonella spp.* harbor over a hundred different T6SS effectors^38,39^. Additionally, each *Salmonella* serovar encodes an individualized set of ∼4 effectors with diverse activities that are hypothesized to facilitate competition with a host-specific microbiota^39^. For example, *Salmonella* Typhimurium encodes a complete T6SS in *Salmonella* pathogenicity island 6 (SPI-6) that is known to harbor at least 4 effector-immunity pairs^40^. These effector-immunity pairs are Tae4-TaiA, Tlde1a-Tldi1a, Rhs1(Tre^Tu^)-RhsI1(Tri^Tu^), and Rhs2-SciX which is also referred to as Rhs^orphan^-RhsI^orphan^. Tae4 is an amidase that is necessary for *Salmonella* Typhimurium to colonize the gut^41–43^, Tlde1a is an L-D transpeptidase that exchanges non-canonical D-amino acids onto the peptidoglycan to promote cell lysis^44,45^, and Rhs1(Tre^Tu^)-RhsI1(Tri^Tu^) is an ADP-ribosyl transferase that targets EF-Tu to inhibit translation^32^. Currently, it has been shown that deletion of Rhs2-SciX in a mouse model decreases virulence and may affect the ability of *Salmonella* Typhimurium to replicate within macrophages^40,46^. Additionally, Rhs2 is an Ntox47 fold and thus a putative nuclease^39^. However, the exact biochemical function and molecular details of the Rhs2-SciX effector-immunity pair are unknown.

To elucidate the molecular details and function of the Rhs2-SciX effector-immunity pair we used a structural approach. We model Rhs2 using AlphaFold to reveal that Rhs2 is structurally homologous to BECR family RNA modifying enzyme. Additionally, we solve an X-ray crystal structure of SciX which reveals a stable globular fold that is also homologous to a BECR family immunity protein. Point-variant toxicity assays in *E. coli* support our structural BECR-family hypotheses for Rhs2-SciX, and allow us to capture Rhs2 bound to EF-Tu. Overall our work expands our knowledge of *Salmonella* effector function and adds to the growing pattern of effectors that target EF-Tu.

## RESULTS

### Rhs2-SciX is an effector-immunity pair

Rhs2 and SciX are predicted to be an effector-immunity pair^40^ because their encoding genes are a in a bicistronic operon within the SPI-6 T6SS of *Salmonella* Typhimurium (Figure 1A). Notably, Rhs2-SciX is encoded adjacent to the effector-immunity pair Rhs1(Tre^Tu^)-RhsI1(Tri^Tu^)^32^. Furthermore, deletion of *rhs2* reduces *S.* Typhimurium virulence and organ dissemination in mouse models suggesting Rhs2 has a toxic activity^40,46^.

**Figure 1:**
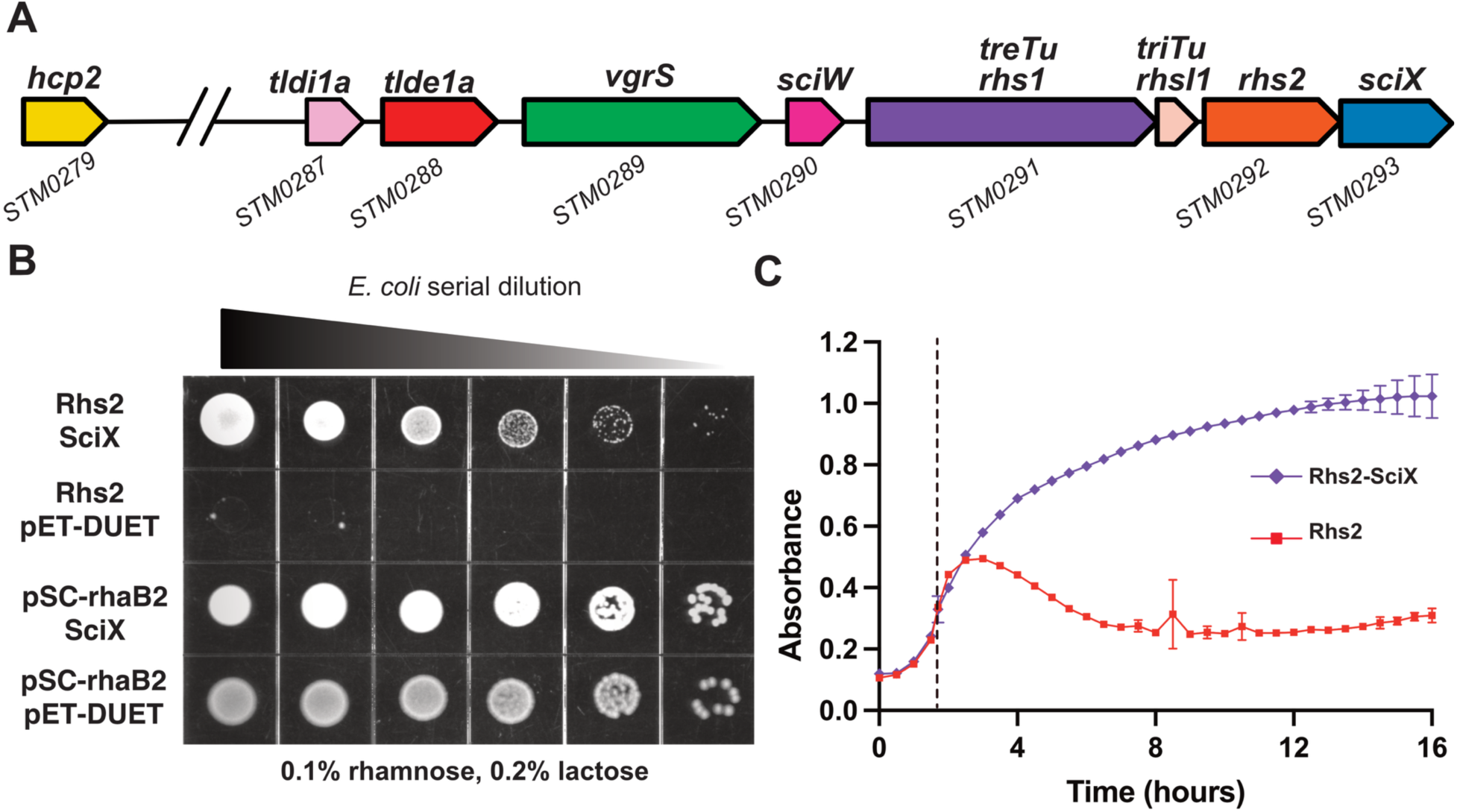
Rhs2-SciX bacterial toxicity and growth assays. **A)** Genomic arrangement of *rhs2* and *sciX* in the T6SS gene cluster of *S.* Typhimurium LT2. Effector-immunity pairs *tlde1a-tldi1a* and *rhs1(treTu)-rhsI1(triTu)* are shown in addition to the spike-tip *vgrS (vgrG)*, the needle protein *hcp2*, and the eag secretion chaperone *sciW*. The *rhsI1* reading frame overlaps with *rhs1*. **B)** Rhs2-SciX effector-immunity toxicity assays by end-point plate. Serial dilutions of *E. coli* co-transformed with *rhs2* pSC-rhaB2 (Rhs2) and *sciX* pETDUET-1 (SciX) or with the empty vectors pSC-rhaB2 and pETDUET-1. **C)** Growth curves of *E. coli* expressing Rhs2 alone (red squares) and the co-expression of Rhs2 and SciX (purple diamonds) induced with 0.1% rhamnose and 0.2% lactose at an of OD600 = 0.3. Point of induction is indicated by dotted line. Samples were run in triplicate.

To test if Rhs2 and SciX are an effector-immunity pair, *rhs2* was placed into the rhamnose inducible plasmid pSC-rhaB2^47^ and *sciX* placed into the lactose inducible plasmid pETDUET-1. After co-transformation of both plasmids into *E. coli* cells, the bacteria were grown and plated by serial dilution onto plates containing rhamnose, lactose, or both inducers (Figure 1B and Figure S1A). Minimal bacterial growth is observed in *E. coli* expressing Rhs2 alone, demonstrating severe toxicity. However, bacterial growth is significantly rescued by cells also expressing SciX (Figure 1B). We did observe that co-transformation of *rhs2* pSC-rhaB2 with *sciX* pETDUET-1 without adding lactose still rescued some bacterial growth (Figure S1A). However, this was likely due to some SciX expression from the T7 promoter even before induction with lactose (Figure S1B).

In parallel with the plate toxicity assays, bacterial growth curves were measured with the induction of Rhs2 and Rhs2-SciX (Figure 1C). Both samples were grown to an optical density at 600 nm (OD_600_) of 0.3 and then induced with both rhamnose and lactose. Induction of Rhs2 expression shows inhibition in growth followed by cell lysis. Additionally, the Rhs2 expressing cells do not recover over the time course of the experiment. Interestingly, induction of Rhs2 was at just under 2hrs but cells continued to grow until the 3hr time point at which point they steadily die over the course of 5hrs. In contrast, cells expressing SciX were protected from Rhs2 toxicity as the cells readily grew to saturation and plateaued (Figure 1C). Overall, our data indicates that Rhs2-SciX can be classified as an effector-immunity pair.

### Rhs2 is a member of the BECR family of nucleases

To determine the mechanism of Rhs2 toxicity we attempted to co-express Rhs2-SciX for structural determination and biochemical characterization. Given the toxicity of T6SS effectors, they are typically expressed bound to their immunity protein. The complex is then denatured and the effector refolded for biochemical assays^31,32,48^. In our hands, soluble Rhs2-SciX complexes for characterization could not be obtained. Instead, we modelled Rhs2 using AlphaFold3^49,50^.

AlphaFold predicts that Rhs2 has a truncated Rhs-cage-like structure N-terminal domain and a globular toxin domain at its C-terminus (Figure 2A and Figure S2). Additionally, the domains are linked by a self-cleavage motif found in Rhs toxins (PxxxxDPxGL)^51–53^. Typically, an Rhs-cage encapsulates the C-terminal toxin domain and self-cleaves after secretion to release the toxin in a prey cell^52,53^. However, compared to other Rhs toxins the Rhs2 cage-like structure is too small to encapsulate the toxin domain making the function of this domain unclear. Despite this, the cleavage motif is highly conserved and still likely separates the cage from the C-terminal toxin domain after secretion. Given this, our structural and functional experiments focused on the toxin domain of Rhs2.

**Figure 2:**
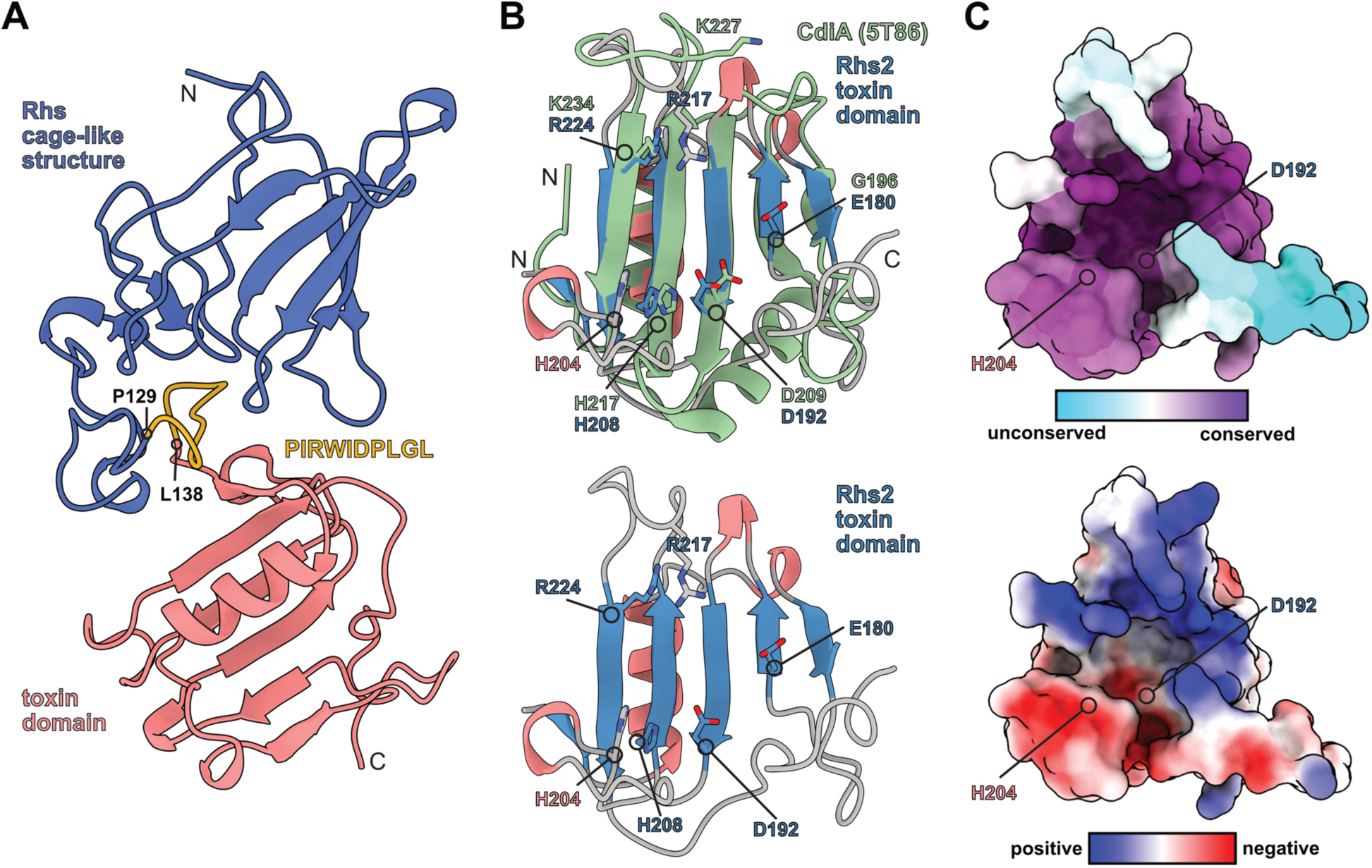
Structural analysis of an Rhs2 model. **A)** AlphaFold3 model of full-length Rhs2. The N-terminal partial Rhs-cage (blue) and the C-terminal toxin (salmon). The domains are linked by the conserved T6SS Rhs-cage cleavage motif (yellow). **B)** Top: Rhs2 toxin domain aligned to an *E. coli* CdiA toxin domain in light green (PDBid 5T86). Putative catalytic residues of BECR family domain RNA modifying enzymes shared with CdiA are shown. Bottom: same view as top of figure, but without overlay with CdiA for clarity. Alpha helices are shown in salmon and beta-strands shown in blue. **C)** Top: surface residue conservation for the Rhs2 toxin domain as calculated by Consurf. Putative catalytic residues D192 and H204 are highlighted. Bottom: relative electrostatic surface potential of Rhs2 calculated by ChimeraX. Putative catalytic residues D192 and H204 are also highlighted.

To gain additional insight into the potential function of Rhs2, we submitted the C-terminal toxin domain AlphaFold3 model to the Dali server^54^. A structure of the toxin CdiA-CT from *E. coli* (PDBid: 5T86) was the top homologue with a Z-score of 7.7 and a root mean square deviation (rmsd) of 3.1 Å (Figure 2B). CdiA is a contact-dependent inhibition (CDI) toxin, which often have nuclease activities^55,56^. Furthermore, CdiA is a member of the BECR fold (barnase/EndoU/colicin E5-D/RelE-like), which is characteristic of many RNases and other RNA modifying enzymes^57^. BECR family members are also known to have the active site residues (D/E/N, H and R/K) to catalyze RNA cleavage^56,57^. When we compare the Rhs2 model with an X-ray crystal structure of CdiA, we clearly observe that Rhs2 contains the BECR family active site residues arranged in a similar conformation to CdiA (Figure 2B). Additionally, residue conservation analysis of Rhs2 using Consurf^58^ demonstrates that the Rhs2 BECR family active site residues are part of a highly conserved surface that creates an active site pocket (Figure 2C top). The surface of Rhs2 is also observed to contain a large positively charged region supporting potential interactions with RNA (Figure 2C bottom). Given our structural analysis, Rhs2 could be a BECR family RNase.

To test our hypothesis that Rhs2 is a BECR family RNase, we created two point variants in the Rhs2 active site (Figure 2B-C). Rhs2 residue D192 was selected as it is at the core of the active site pocket, identical in position to CdiA residue D209, and a potential nucleophile for RNA cleavage. Rhs2 residue H204 was chosen because it is on a conserved loop that projects from the active site that is in a different predicted conformation from CdiA (Figure 2B). As histidine residues are frequently part of an RNase motif^57^ this could indicate a unique molecular detail in the mechanism of Rhs2 compared to other BECR folds. Both residues were substituted with alanine (D192A and H204A). Using the Rhs2 point variants, we repeated *E. coli* toxicity assays to test whether these residues are important for the biochemical function of Rhs2. Both point variants completely abrogate Rhs2 toxicity as observed by end-point plate assays (Figure 3A) and bacterial growth curves (Figure 3B). These data indicate that these residues likely participate in the nuclease activity of Rhs2 and/or are necessary for substrate binding. Taken together with our structural analysis we conclude that Rhs2 is a BECR family nuclease.

**Figure 3:**
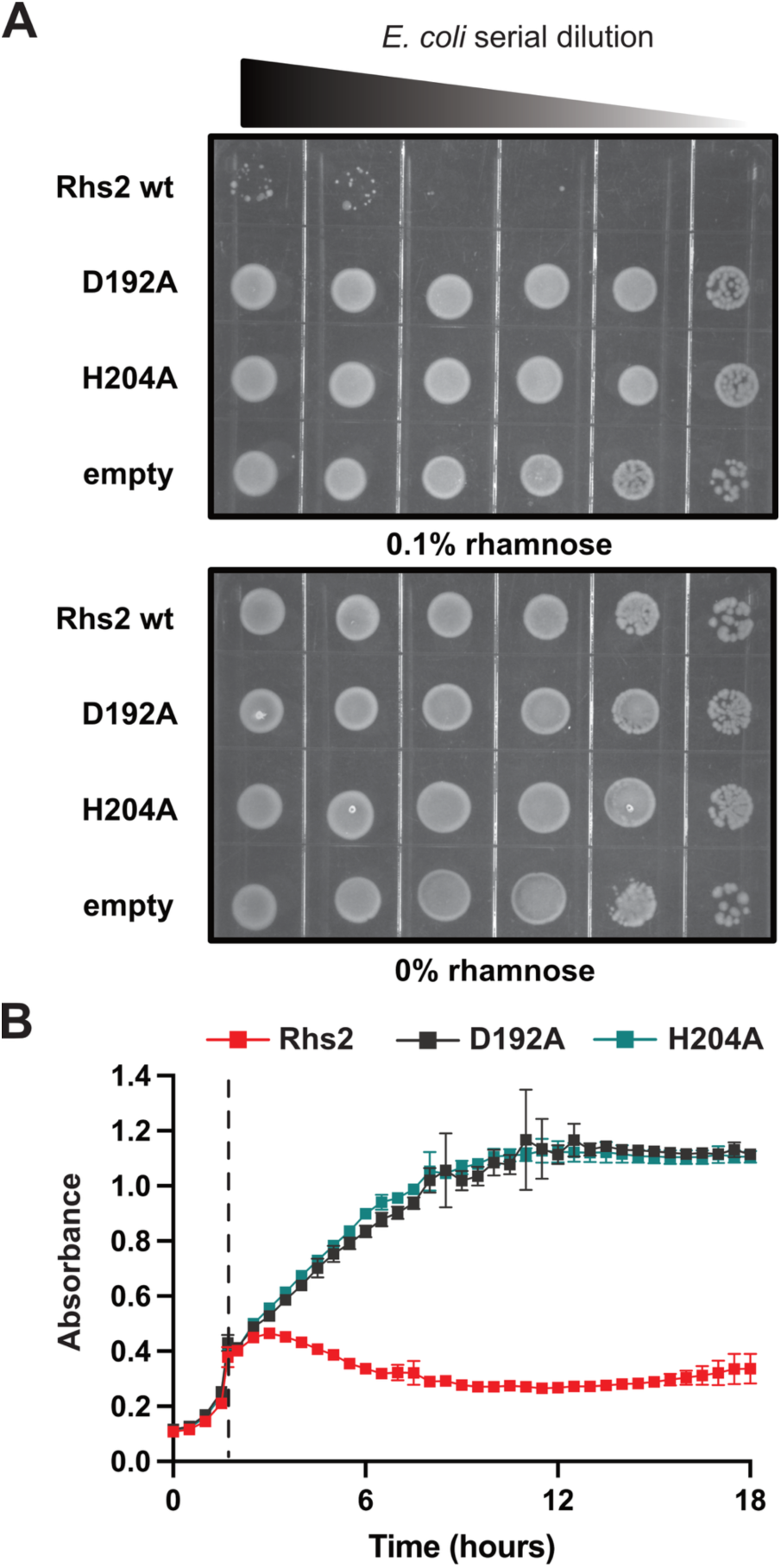
Rhs2 variant bacterial toxicity and growth assays. **A)** Rhs2 and point variant toxicity assays by end-point plate. Serial dilutions of *E. coli* transformed with pSC-rhaB2 containing wild-type Rhs2 (Rhs2 wt), Rhs2 active site point variant D192A, Rhs2 active site point variant H204A, and pSC-rhaB2 empty. Top panel is grown with 0.1% rhamnose to induce Rhs2 expression and the bottom panel lacks rhamnose and Rhs2 expression. **B)** Growth curves of *E. coli* expressing Rhs2 wild-type (red squares), Rhs2 D192A (black squares), and Rhs2 H204A (blue-green squares). Samples were induced with 0.1% rhamnose at an of OD600 = 0.3. Point of induction is indicated by dotted line. Samples were run in triplicate.

### Biophysical analysis of SciX reveals a conserved surface with limited flexibility

Although we could not obtain a Rhs2-SciX complex for structural studies, SciX alone was readily purifiable. After buffer optimization using nano differential scanning fluorimetry (nanoDSF) (Figure S3A) we were able to obtain an X-ray crystal structure of SciX (Figure 4A and Table 1). It is important to note that crystals were only obtained after an incubator failure during a power outage that heated then re-cooled the crystallization trials to 4°C. Additionally, SciX crystallized with two nearly identical copies in the asymmetric unit although SEC-MALS analysis indicates that SciX is a monomer in solution (Figure S3B, Figure S4, Table S1).

**Figure 4:**
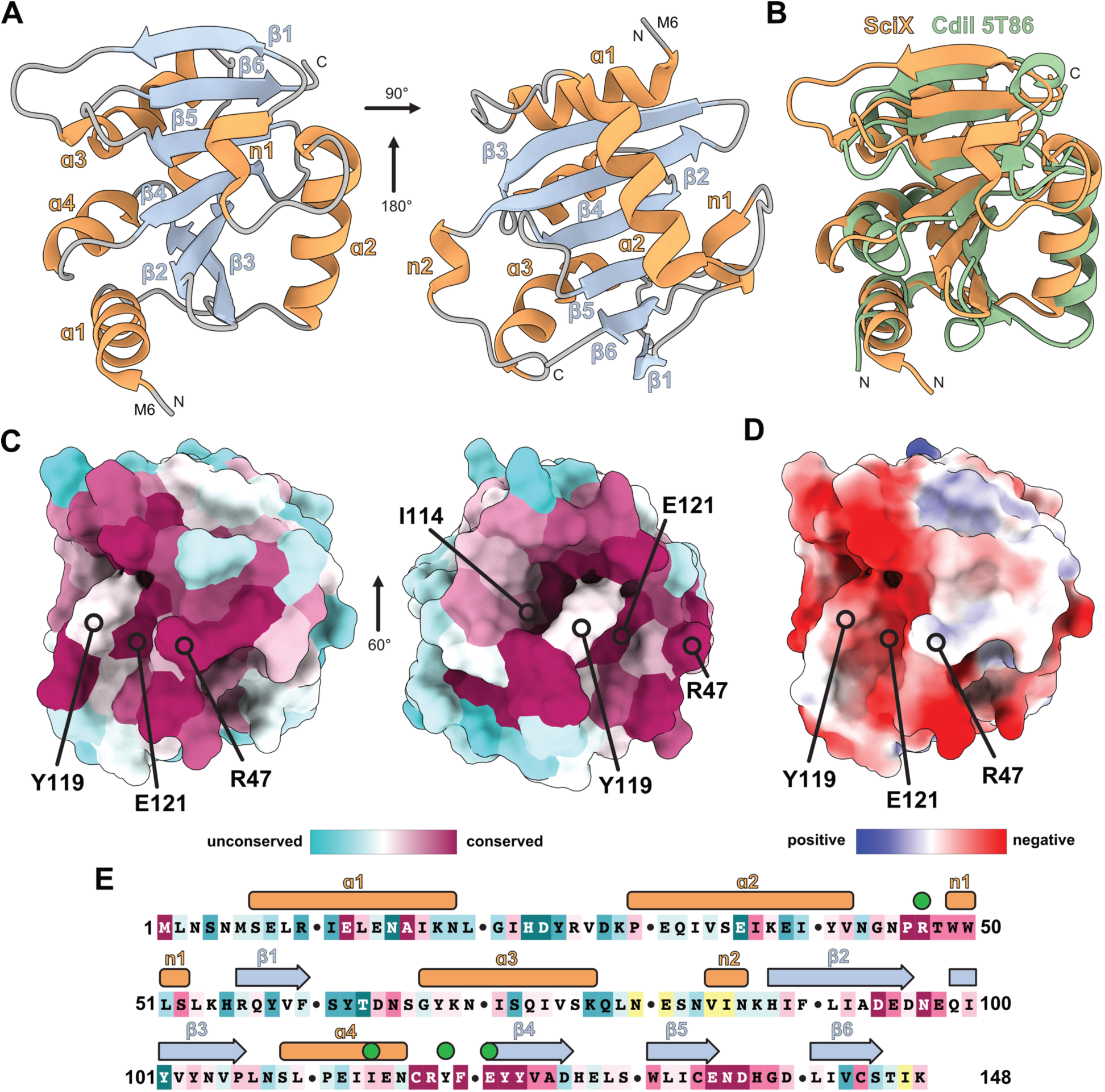
An X-ray crystal structure of SciX. **A)** X-ray structure of SciX annotated by secondary structure (PDBid 9ZR2). Two views of SciX are shown relative to each other by the indicated rotations. Alpha helices are shown in light orange and beta-sheets are shown in light blue. **B)** Structural alignment of SciX and CdiI (PDBid 5T86). CdiI is in the immunity protein for CdiA. SciX is shown in orange and CdiI in light green. **C)** Surface residue conservation of SciX as calculated by Consurf. Two views of the conserved surface of SciX are shown by the indicated rotation. Residues at the core of the conserved surface are indicated and labeled. **D)** Electrostatic surface potential of the SciX conserved surface calculated by ChimeraX and the adaptive Poisson-Boltzman solver. Residues at the core of the conserved surface are indicated and labeled. **E)** Sequence of SciX annotated by residue conservation as calculated in panel C and the observed secondary structure from the crystal structure. Residues highlighted in panel C are indicated by green circles.

**Table 1:** Data collection and refinement statistics for SciX.

|  |  |
| --- | --- |
| **Data Collection** | SciX |
| Wavelength (Å) | 0.92 |
| Space group | P 1 21 1 |
| Cell dimensions |  |
| a, b, c (Å) | 41.07 83.63 41.63 |
| $\alpha, \beta, \gamma$ (°) | 90 99.68 90 |
| Resolution (Å) | 41.82 - 2.3 (2.38 - 2.3) |
| Total reflections | 822205 (7414) |
| Unique reflections | 12338 (1202) |
| CC (1/2) | 1.000 (0.92) |
| $R_{merge}$ | 0.05 (0.35) |
| $R_{pim}$ | 0.03 (0.23) |
| $I/\sigma$ | 19.2 (4.3) |
| Completeness (%) | 99.57 (97.0) |
| Redundancy | 6.7 (6.4) |
| **Refinement** |  |
| $R_{work} / R_{free}$ (%) | 22.7 / 25.5 |
| Average B-factors | 51.81 |
| Protein | 57.96 |
| Water | 47.38 |
| No. atoms |  |
| Protein | 2360 |
| Water | 33 |
| Rms deviations |  |
| Bond lengths (Å) | 0.002 |
| Bonds angles (°) | 0.59 |
| Ramachandran plot (%) |  |
| Total favoured | 95.67 |
| Total allowed | 3.61 |
| PDB code | 9ZR2 |
| Statistics for the highest resolution shell are shown in parenthesis |  |

The structure of SciX consists of a central 6 strand beta-sheet (β1-β6) flanked by 4 alpha helices (ɑ1-ɑ4) and two 3_10_ helices (n1 and n2) (Figure 4A). When the structure of SciX was submitted to the Dali server, the top structural homologue was CdiI (PDBid 5T86) with a Z-score of 9.9 and a rmsd of 2.7 Å (Figure 4B). Importantly, CdiI is the immunity protein for CdiA which is also the structural homologue of Rhs2 (Figure 2B). The surface of SciX was also analyzed for residue conservation using Consurf^58^, and for electrostatics using both ChimeraX^59^ and the adaptive Poisson-Boltzmann solver^60^ (Figure 4C-D). SciX contains a conserved surface that primarily consists of the connecting loops between the core beta-sheet and the n1 (3_10_ helix) between ɑ2 and β1. Furthermore, the SciX conserved surface is negatively charged suggesting electrostatic complementation with part of the Rhs2 active site surface (Figure 4D and Figure 2C).

A detailed look at the putative Rhs2 binding site on SciX highlights 4 residues of interest (Figure 4C and Figure 4E). Specifically, conserved residues R47, E121, and I114 are interrupted by the variable residue Y119. Because T6SS effectors have subtle molecular modifications that dictate new toxic functions^31,45^, their immunity proteins must also have unique structural features to bind their cognate effector^10^. Given this, it is tempting to speculate that because Y119 is not conserved in the CdiI BECR toxin-immunity fold it may represent binding specificity to Rhs2.

As crystals were not obtained until an incubator failure, we had collected a triple-resonance NMR data set to study the solution state structure of SciX. The assigned ^1^H-^15^N HSQC of SciX is shown in Figure 5A. Clear peak dispersion indicates that in solution SciX is a well folded globular protein. Additionally, the calculated solution-state secondary structure from the backbone assignments (Cɑ, Cβ, CO, and H-N) using TALOS^61,62^ closely matches the secondary structure observed in the crystal structure (Figure 5B). We also measured SciX protein dynamics by collecting a ^1^H-^15^N NOE heteronuclear relaxation experiment (Figure 5B). A ^1^H-^15^N NOE heteronuclear relaxation experiment allows visualization of which regions adopt a stable conformation (NOE ≥ 0.6) and which regions are dynamic (NOE<0.6 or negative NOE if disordered)^63,64^. The data shows that aside from the cloning artifact, SciX is predominantly globular and rigid at least in our buffer conditions. We do highlight that several of the loop regions could not be assigned or produced no detectable signal in the ^1^H-^15^N NOE relaxation experiment (no NOE bar). A lack of signal is typical of intermediate exchange where the loop regions are changing conformation on the time scale of the NMR experiment^65,66^. Interestingly this also included most of beta-strand β5 suggesting this region of the SciX structure is dynamic. Overall, the NMR data demonstrates that the solved crystal structure also represents the solution state structure of SciX, and that SciX is primarily a rigid globular fold.

**Figure 5:**
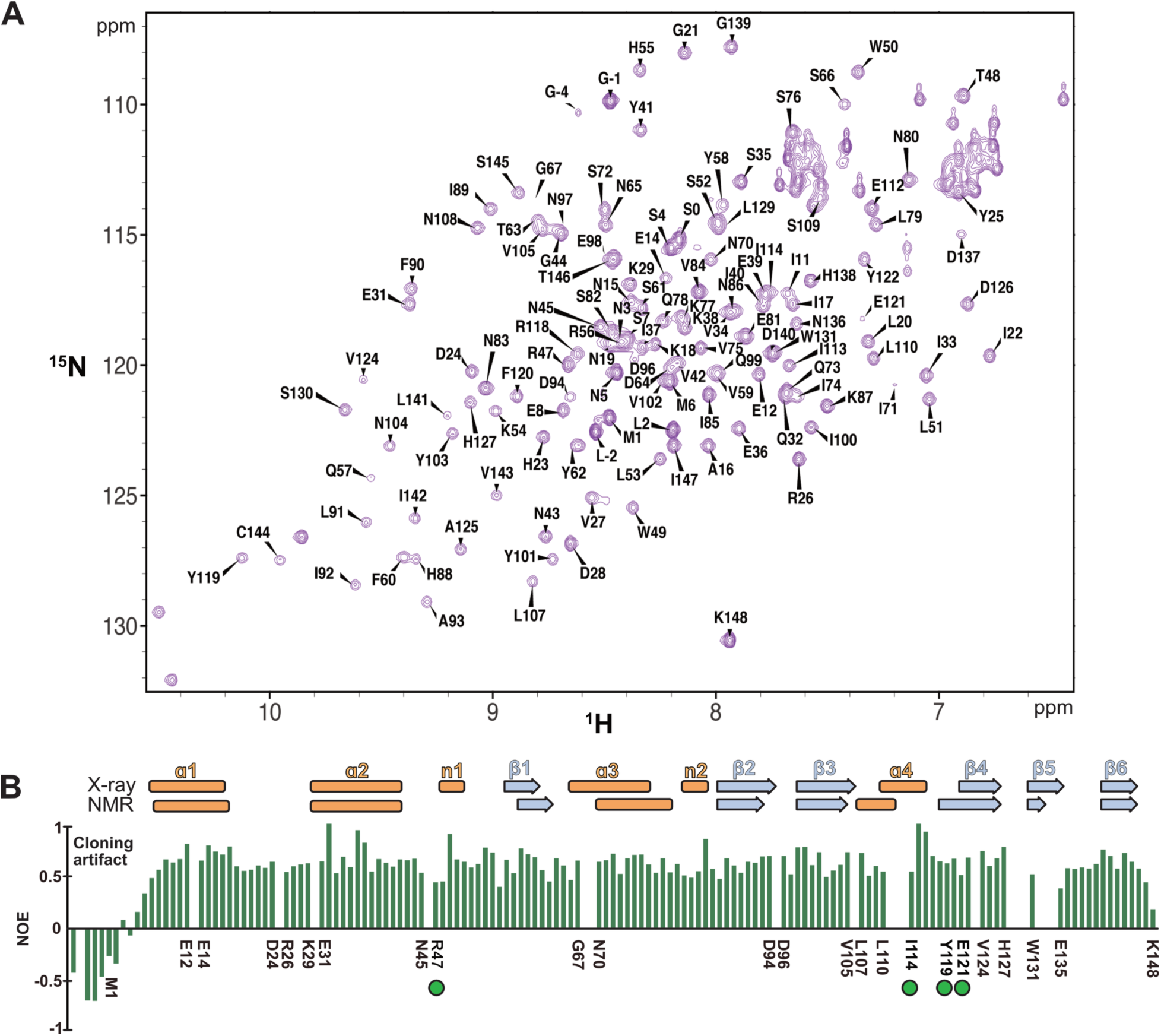
Solution state structure and dynamics of SciX. **A)** Assigned ^1^H-1^5^N HSQC spectra of SciX. **B)** top: secondary structure of SciX determined by X-ray crystallography as compared to calculated by assigned backbone chemical shifts. Alpha-helices are shown in light orange and beta-strands in light blue. Bottom: ^1^H-^15^N heteronuclear NOE relaxation data for SciX plotted by residue and aligned to the secondary structure annotation. NOE peak height is shown by green bars. Residues at the boundaries of dynamic regions are indicated. SciX residues on the conserved surface highlighted in Figure 4 are also indicated by green circles.

### Molecular model of a Rhs2-SciX complex

To help validate the predicted Rhs2 active site (Figure 2) and the Rhs2 binding surface of SciX (Figure 4), we also used AlphaFold3^50^ to model a Rhs2-SciX complex (Figure 6 and Figure S5). Overall AlphaFold3 predicts that the conserved binding surface on SciX binds directly to the Rhs2 active site. Additionally, the configuration of the Rhs2-SciX complex is nearly identical to the CdiA-CdiI complex (PDBid 5T86) (Figure 6A). Upon examining the predicted Rhs2-SciX interface, SciX appears to make contacts with the BECR family active site residues of Rhs2. SciX R47 likely makes an H-bond or salt bridge with Rhs2 D192 and SciX H138 appears to contact both Rhs2 H204 and H208 (Figure 6B). These contacts are less than 4 Å in distance and many are near 3.5 Å which is likely within the error of the AlphaFold prediction^67^. Importantly, both Rhs2 D192 and H204 are required for toxicity (Figure 3) and buried in the Rhs2-SciX interface (Figure 6). Additionally, SciX residue Y119 packs against Rhs2 residue R224 which could position R224 to help the residue form a salt bridge with SciX E121 (Figure 6C). As Y119 is unconserved (Figure 4C and 4E) this could be unique to SciX recognition and binding of Rhs2. Finally, the side chain of SciX I114 packs against SciX F120 to stabilize the loop containing both residues SciX Y119 and E121.

**Figure 6:**
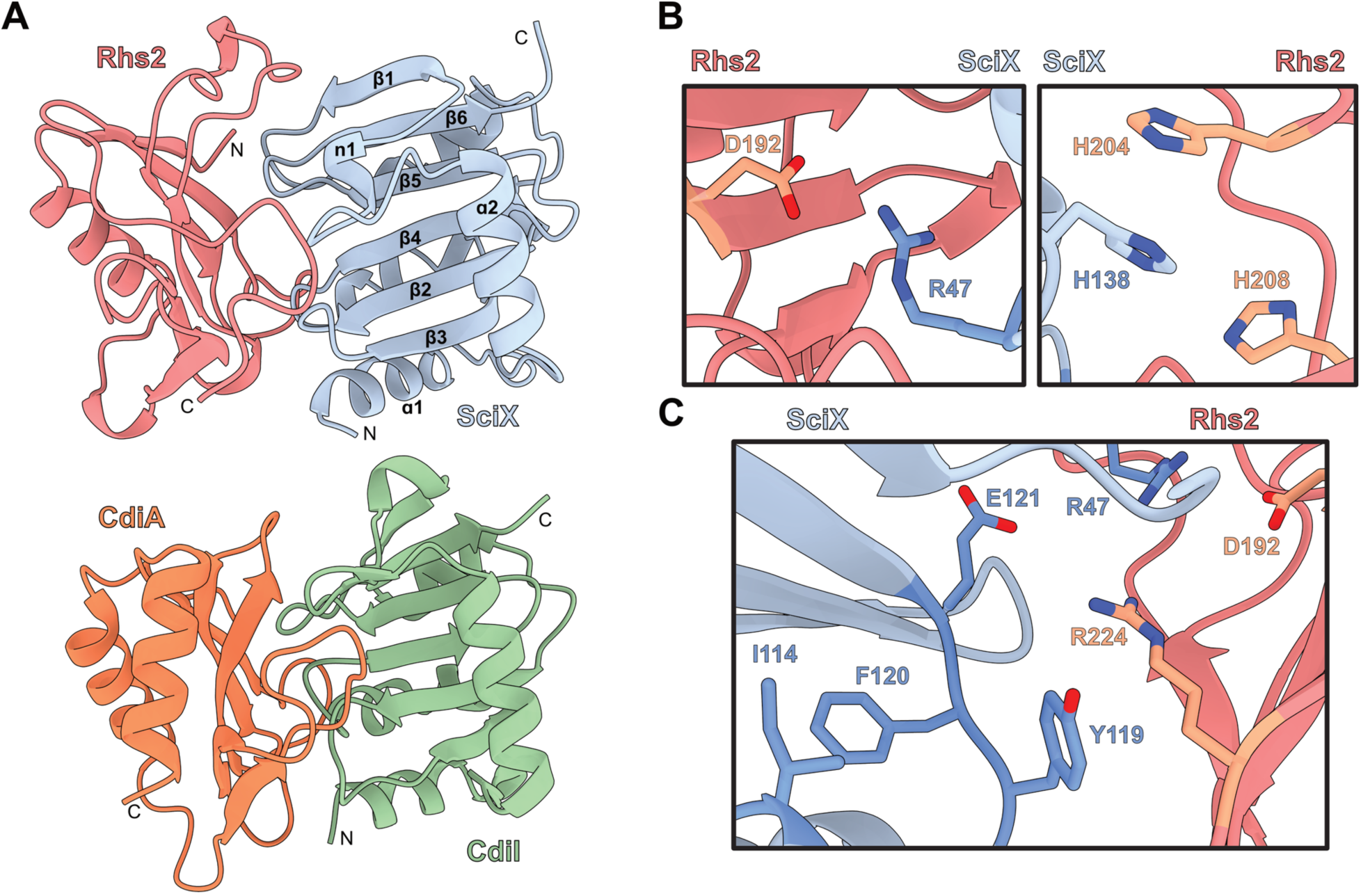
Predicted model of an Rhs2-SciX complex. **A)** AlphaFold model of Rhs2-SciX compared to CdiA-CdiI (PDBid 5T86). Rhs2 is shown in salmon, SciX in light blue, CdiA in bright orange, and CdiI in light green. **B)** Predicted residue interactions between the Rhs2 active site surface and the conserved SciX surface. Rhs2 D192 may contact SciX R47 and SciX H138 may interact with both Rhs2 H204 and H208. **C)** SciX Y119 may pack against Rhs2 R224 to position the residue for interaction with SciX E121. The SciX structure demonstrates that I114 packs against F120 to stabilize the loop conformation for Y119 and E121.

To test if the highlighted SciX residues are important for binding and inhibition of Rhs2 toxicity, each SciX residue shown in Figure 6 (R47, I114, Y119, F120 and E121) was substituted with alanine and cloned into pETDUET-1. The SciX point variants were then co-transformed with Rhs2 and toxicity assays performed in *E. coli* by both plate assay and growth kinetics. Figure 7 demonstrates that each individual residue does not contribute significantly to SciX inhibition of Rhs2. This was observed by both plate spotting assays (Figure 7A-C) and by growth curve kinetics (Figure 7D). Each individual variant growth assay is also shown in Figure S6 for clarity. These results could infer that there is significant redundancy in the SciX binding interface to assure inhibition of Rhs2. Namely that a single disrupted interaction is insufficient to disrupt Rhs2 binding. Alternatively, the AlphaFold modelled Rhs2-SciX interface is incorrect. However, given the high structural and functional similarity to the CdiA-CdiI complex, the overall AlphaFold3 model is likely correct although the specific residue contacts have error.

**Figure 7:**
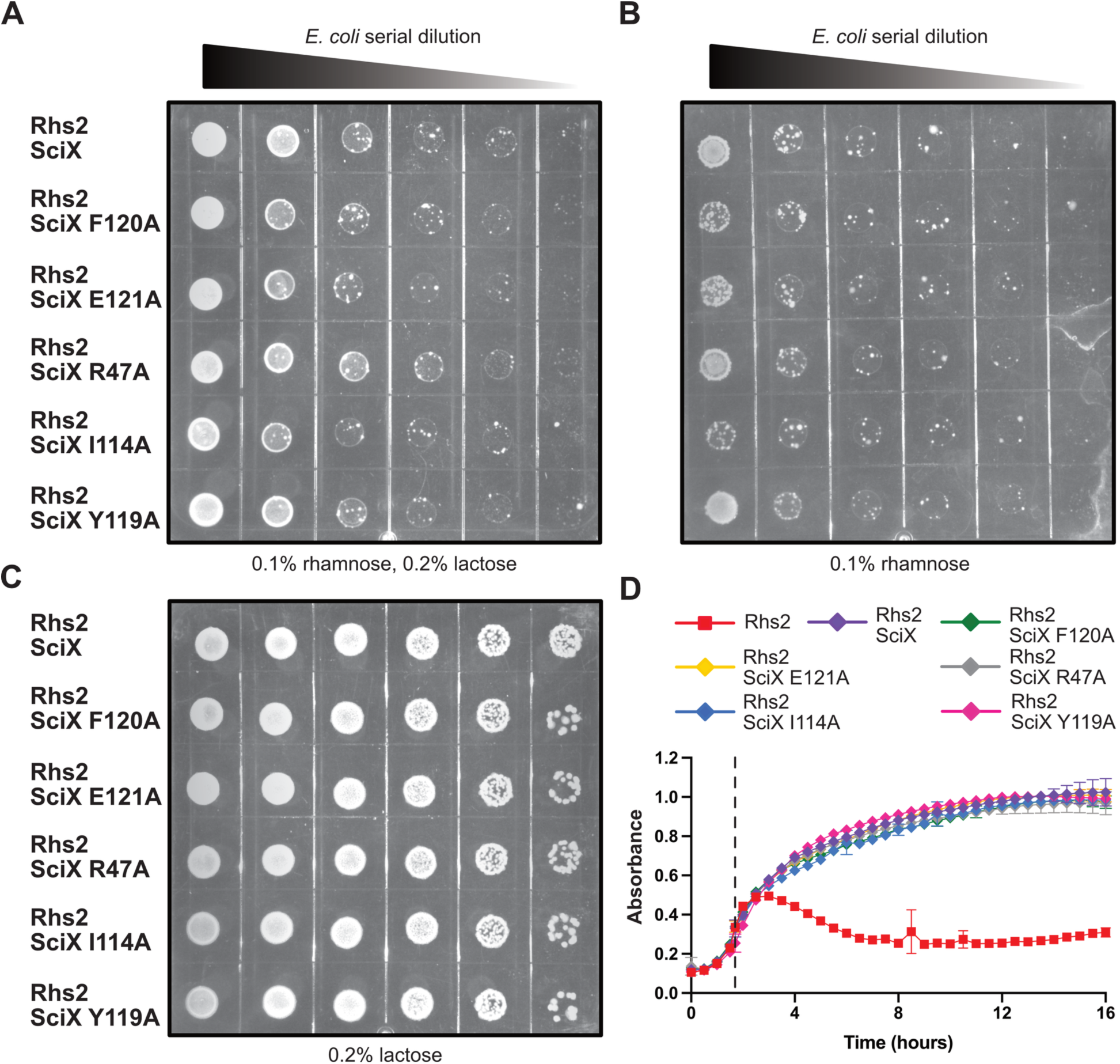
Rhs2 and SciX variant bacterial toxicity and growth assays. **A)** Rhs2 and SciX point variant toxicity assays by end-point plate. Serial dilutions of *E. coli* co-transformed with *rhs2* pSC-rhaB2 (Rhs2) and *sciX* pETDUET1 (SciX) or with a SciX point variant in pETDUET1. Plates contained 0.1% rhamnose and 0.2% lactose to induce both Rhs2 and SciX. **B)** Same assay as panel A, but containing 0.1% rhamnose to induce only Rhs2. **C)** Same assay as panel A and B, but containing 0.2% lactose to induce only SciX. **D)** Growth curves of *E. coli* expressing Rhs2 wild-type (red squares) or co-expressed with wild-type SciX and each SciX variant indicated by color. Samples were induced with 0.1% rhamnose and 0.2% lactose at an of OD600 = 0.3. Point of induction is indicated by dotted line. The Rhs2 curve is repeated from previous figures. Samples were run in triplicate.

### Rhs2 binds directly to EF-Tu

Since we lacked an experimental Rhs2 structure, we attempted to purify the catalytically inactive Rhs2 variants for both structure determination and binding assays with SciX. Based on the Rhs2-SciX variant toxicity assays, a single point variant should not disrupt complex formation. Given this, a purified Rhs2 variant is likely usable for ^1^H-^15^N HSQC NMR titration assays with ^15^N labeled SciX to determine the Rhs2 binding interface^64,68^. The minimal toxin domain of Rhs2 (residues 137-246) with the point variant D192A or H204A were cloned with a C-terminal 6His-tag in pET29b+, recombinantly expressed, and purified by affinity chromatography. Both variants co-purified with a larger molecular weight protein of close to 45 kDa (Figure 8A and Figure S7). To determine the identity of the co-purified protein, the 45 kDa band was cut from the gel and the material identified by mass-spectrometry (LC-MS/MS). The mass-spectrometry results unequivocally identified the protein as EF-Tu from *E. coli* (Table 2). EF-Tu functions in protein synthesis by binding amino-acylated tRNAs and piloting them to a translating ribosome^69^.

**Figure 8:**
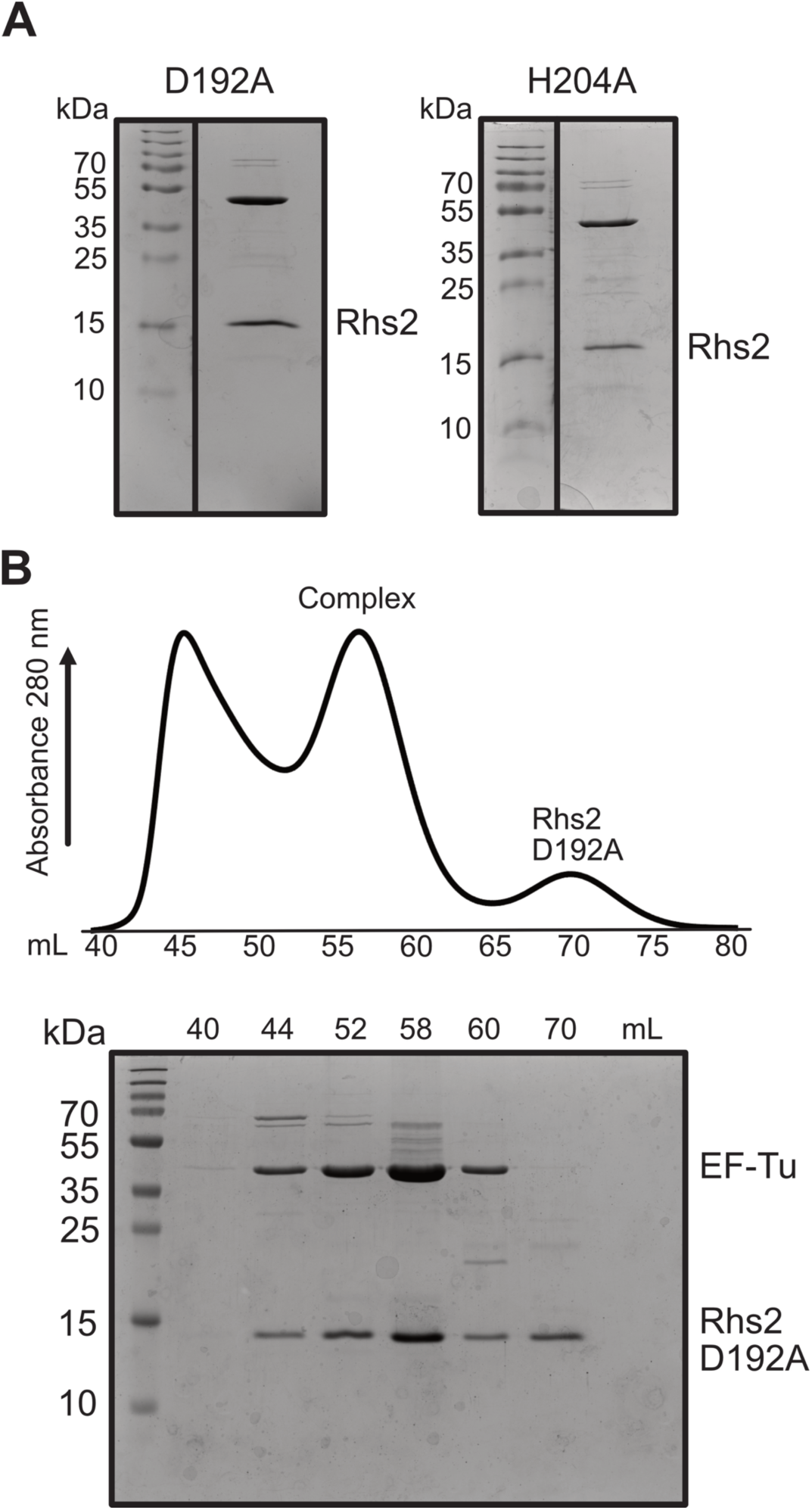
Purification of Rhs2 variants bound to EF-Tu. **A)** Coomassie stained SDS-PAGE gels of 6His-tagged Rhs2 toxin domain variants D192A and H204A purified by metal affinity chromatography. The Rhs2 toxin domain (residues 137-246) is labeled. **B)** Size exclusion chromatography (SD75 HiLoad 16/600) trace of purified Rhs2 D192A and EF-Tu. The complex peak is indicated in addition to Rhs2 D192A alone. A Coomassie stained SDS-PAGE gel of select fractions is shown with EF-Tu and Rhs2 D192A labeled.

**Table 2:** Rhs1 protein binding partner identification by mass-spectrometry.

| **Accession** | **Gene** | **Organism** | **Protein** | **Coverage (%)** | **Peptides** | **Counts** |
| --- | --- | --- | --- | --- | --- | --- |
| P0A1H5 | *tufA* | *E. coli* | Elongation factor Tu | 97.97 | 63 | 2516 |
| P0CE48 | *tufB* | *E. coli* | Elongation factor Tu 2 | 97.97 | 63 | 2515 |
| P23908 | *argE* | *E. coli* | Acetylornithine deacetylase | 57.96 | 18 | 51 |
| P77434 | *alaC* | *E. coli* | aminotransferase AlaC | 62.86 | 24 | 45 |
| P0A1R0 | *hfq* | *S. Typhimurium* | RNA-binding protein Hfq | 38.24 | 6 | 9 |
| P77581 | *astC* | *E. coli* | Succinylornithine transaminase | 58.62 | 17 | 25 |
| P77302 | *dgcM* | *E. coli* | Diguanylate cyclase DgcM | 30.49 | 11 | 16 |
| P0ABH0 | *ftsA* | *E. coli* | Cell division protein FtsA | 42.86 | 11 | 15 |
| P06987 | *hisB* | *E. coli* | Histidine biosynthesis HisB | 38.03 | 11 | 11 |
| P0A6B7 | *iscS* | *E. coli* | Cysteine desulfurase IscS | 32.18 | 12 | 12 |

Binding partner hits are listed by Uniprot accession number, gene name, organism and functional annotation. Coverage refers to percentage of expected protein amino acid sequence identified, peptides is the number of unique peptide sequences measured, and counts is the total spectra assigned or relative abundance of each protein.

We next purified an Rhs2-EF-Tu variant complex by both metal affinity chromatography followed by size-exclusion chromatography (SEC). As shown in Figure 8B, Rhs2 D192A forms a direct interaction with EF-Tu and remains a stable complex as assayed by SEC. However, we did observe that Rhs2 D192A may dissociate from EF-Tu or Rhs2 maybe saturated relative to cellular EF-Tu levels (Figure 8B elution volume 70 mL). Additionally, the variant complex does form aggregates indicating that it might be unstable (Figure 8B elution volume 44 mL). In line with this observation, Rhs2 H204A precipitated readily after affinity chromatography and could not be used for SEC. Together, this shows that the residues D192 and H204 are not required for EF-Tu binding and that the active site of Rhs2 may be exposed to solvent when in complex with EF-Tu. The Rhs2 variant EF-Tu complexes were of low yield and prone to precipitation, making structural determination difficult. Significant construct optimization remains to acquire an experimental Rhs2-EF-Tu complex structure, or to obtain enough Rhs2 material to separate from EF-Tu and refold for NMR titration experiments with SciX. Regardless our data demonstrates that Rhs2 binds directly to EF-Tu. Importantly, EF-Tu binding is observed frequently by T6SS^32,33,70^ and CDI toxins^57,71^ and often results in cell death by inhibiting protein translation. Given this, Rhs2 also likely functions by disrupting a step in the translation process.

## DISCUSSION

Here we show that the T6SS effector-immunity pair Rhs2-SciX is structurally homologous to a CdiA-CdiI contact-dependent toxin-immunity pair from *E. coli.* Additionally, an AlphaFold model of Rhs2 combined with Rhs2 point variant toxicity assays in *E. coli* demonstrate that structurally Rhs2 is member of the BECR family of nucleases. We also solve an X-ray structure of SciX, probe its solution state dynamics, and compare the SciX structure with CdiA-CdiI. Together, this supports the conclusion that SciX inhibits Rhs2 by active site occlusion and contacts key Rhs2 active site residues. Finally, we discover that Rhs2 binds directly to EF-Tu. As Rhs2 is a BECR fold that binds EF-Tu, we propose that its toxicity is likely due to modifying RNA to inhibit translation.

T6SS effectors have been shown to directly target RNA for toxicity. These effectors include RhsP2 from *Pseudomonas aeruginosa* that ADP-ribosylates double stranded RNA^31^, and TseR that from *Yersinia pseudotuberculosis* that functions as an RNase^72^. Given their substrates, both effectors cause cell death by the inhibition of translation and are trans-kingdom effectors that kill both prokaryotic and eukaryotic cells. However, RhsP2 is an ART-fold (ADP-ribosyltransferase) that is structurally unrelated to Rhs2^31^ and TseR has no experimental structural data available for comparison with Rhs2. In contrast, the AlphaFold model of Rhs2 was found to be structural similar to contact-dependent toxins that contain BECR domains (Figure 2). Specifically, an AlphaFold3 model of Rhs2 and an X-ray crystal structure of SciX are homologous to the toxin-immunity protein pair CdiA-CdiI from *E. coli* (PDBid: 5T86). Although no publication describing a mechanistic characterization is associated with the PDB entry 5T86, the CDI complex did allow the classification of Rhs2 as a BECR fold. Importantly, Rhs2 has a catalytic active site configuration consistent with a BECR family RNase (D/E/N, H and R/K)^57^. For Rhs2 this includes residues D192 and H204 that are required for toxicity (Figure 2–3).

Direct binding to EF-Tu is observed frequently with both T6SS effectors^32,33,70^ and CDI toxins^57,71^. For T6SS effectors, binding to EF-Tu can be used to facilitate membrane translocation by prePAAR effectors such as Tse6 in *Pseudomonas aeruginosa*^33^ and Rhs2 from *Serratia marcescens*^70^. Note that Rhs2 from *Serratia marcescens* is a DNase that is unrelated to Rhs2 from *Salmonella* Typhimurium. Alternatively other T6SS effectors such as Tre^Tu^ (Rhs1) from *S.* Typhimurium directly modify EF-Tu to inhibit translation^32^. For CDI toxins, many have been shown to bind EF-Tu to pilot them to their RNA substrates. CdiA from enterohemorrhagic *E. coli* EC869 and CdiA from *Klebsiella pneumoniae 342* both bind EF-Tu, which is essential for their tRNA cleavage activities^57,73^. Since Rhs2 is a BECR family fold enzyme with high structural similarity to CdiA toxins and binds directly to EF-Tu, we hypothesize that Rhs2 cleaves RNA to inhibit translation. Additionally, based on the structural and functional similarities to CdiA toxins, the most likely substrate for Rhs2 is tRNA. To prove this hypothesis, recombinant Rhs2 would need to be purified for *in vitro* activity assays and/or whole-cell RNA isolated and analyzed by mass-spectrometry. However, to date we have been unable to isolate purified wild-type Rhs2 for assays or harvest enough *E. coli* for RNA-based assays due to the toxicity of Rhs2 (Figure 2).

To our knowledge, very few if any T6SS immunity protein structures have been solved without being in complex with their cognate effector. Here, were solve a structure of the immunity protein SciX alone and characterize its solution state properties by NMR spectroscopy (Figure 4 and Figure 5). Our biophysical data demonstrates that SciX is a monomeric globular fold that shows very little conformational dynamics in solution. Specifically, the heteronuclear NOE relaxation data demonstrate that only the short loop regions between secondary structure elements are flexible and are undergoing movement in solution. Taken together with the Rhs2-SciX AlphaFold3 model, our structural and dynamic data suggests that SciX is rigid and undergoes little to no conformational change upon binding Rhs2. Namely, SciX has co-evolved to rigidly compliment the structure of Rhs2 and socket into the Rhs2 active site to inhibit toxicity.

Previous studies have shown that Rhs2 is required for pathogenicity in a mouse model^40^ and contributes to *Salmonella* replication in macrophages^46^. However, unlike Rhs1 (Tre^Tu^), Rhs2 lacks a full Rhs-cage and PAAR domain for T6SS secretion. This is in part why Rhs2 is referred to as Rhs^orphan^. For Rhs1, the N-terminal region contains a prePAAR motif and PAAR domain for loading onto VgrS (VgrG)^74^ which requires the <u>e</u>ffector <u>a</u>ssociated <u>g</u>ene (Eag) chaperone SciW^24,75^. SciW also binds the transmembrane domain (TMD) of Rhs1 during T6SS loading. After secretion, the TMD is responsible for helping translocate Rhs1 across the prey cell inner membrane into the cytoplasm where EF-Tu is located^24,75^. If Rhs2 lacks both a PAAR domain and a TMD, this raises the question of how Rhs2 is both secreted by the T6SS and subsequently translocated into the cytoplasm to bind EF-Tu and degrade any nucleic acid substrate. The first possibility is that Rhs2 requires a yet to be discovered adaptor that allows it to be loaded into the lumen of one of the three *Salmonella* Hcps^43,76^ or facilitates binding to a structural element on VgrS^74^. However, this still would not explain how Rhs2 reaches the cytoplasm of a prey cell. Another possibility is that the partial cage of Rhs2 binds to the C-terminal region cage of Rhs1 and extends the structure. An Rhs cage is made of Y-D residue repeats that assemble a hollow curved beta-barrel structure^35,52^. It is possible that the Y-D repeat regions of Rhs2 assemble relative to an N or C-terminal beta-strand where one side does not have its backbone hydrogen bonding satisfied and extend the cage to include Rhs2. This of course does not solve the lack of TMDs for inner membrane translocation. In this scenario, Rhs2 would have to “piggyback” on the TMD mediated translocation of Rhs1. Instead, Rhs2 may recognize a conserved Gram-negative inner membrane protein to help facilitate translocation into the cytoplasm. Regardless, significant work remains to elucidate the secretion mechanism of Rhs2.

Our work has shown that Rhs2-SciX are a bonafide T6SS effector-immunity pair, and that Rhs2 is a BECR family nuclease that likely inhibits translation to elicit cell death. Overall, these findings have advanced our understanding of *Salmonella* T6SS effectors and added to the list of effectors that utilize the essential protein EF-Tu for toxicity.

## MATERIALS AND METHODS

### Design of protein expression and gene constructs

A protein expression construct of SciX or STM0293 (*S.* Typhimurium LT2) was synthesized and cloned into pET-21a+ with a cleavable N-terminal 6His-tag for purification purposes (Genscript). For toxicity experiments, SciX was also cloned into multiple cloning site 2 of pETDUET-1 tagless with restriction sites NdeI/XhoI to be co-transformed with Rhs2 (Genscript). Point variants of SciX were achieved using Q5 mutagenesis (New England Biolabs). Rhs2 or STM0292 was first cloned in pET21a+ with sites NdeI/XhoI (Genscript) then subcloned into pSC-rhaB2 with the sites NdeI/NcoI. Rhs2 point variants D192A and H204A were synthesized in pET29b+ (Twist Bioscience) using the restriction sites NdeI/XhoI then subcloned into pSC-rhaB2 with the sites NdeI/HindIII. For recombinant protein expression, the minimal toxin domain of Rhs2 (residues 137-246, Cterm) was synthesized with point variants D192A and H204A then cloned into pET29b+ using restrictions sites NdeI/XhoI to contain a C-term 6His-tag (Twist Bioscience). The synthesized point variants were subcloned into pSC-rhaB2. All clones were transformed into *E. coli* BL21(DE3)Gold competent cells for protein expression and toxicity assays.

### Bacterial toxicity assays

Wild-type Rhs2 (pSC-rhaB2) and SciX (pETDUET-1) wild-type and SciX point variants were co-transformed into BL21(DE3)Gold electrocompetent cells via electroporation. Controls included the co-transformation with cognate empty vectors. Cultures were grown at 37°C O/N, diluted to an OD600 of 0.6 the following day, then further serially diluted from 10^-1^ to 10^-5^. 5 μL of each sample was spotted onto three different Lysogeny Broth (LB) agar plates: 0.1% rhamnose alone, 0.2% lactose alone and 0.1% rhamnose/0.2% lactose. Plates were grown at 37°C then visualized after 24hrs. All plates for Rhs2-SciX experiments contained 100 μg/mL ampicillin (pETDUET-1) and 50 μg/mL of trimethoprim (pSC-rhaB2). For Rhs2 wild-type and point-variant toxicity assays, the plates only contained 50 μg/mL of trimethoprim with or without 0.1% rhamnose.

### Bacterial growth assays

Cultures of *E. coli* BL21(DE3)gold containing pSC-rhaB2 and/or pETDUET-1 with and without Rhs2 and SciX wild-type or variants were grown O/N at 37°C with the appropriate antibiotics (50 μg/mL trimethoprim for pSC-rhaB2, 100 μg/mL ampicillin for pETDUET-1, and both antibiotics for co-transformations). The following day, cultures were diluted to an OD600 nm of 0.1 in a total of 150 μL. Cultures were added to 96 well plates for measuring kinetic growth curves using a Synergy HTX multi-mode plate reader (BioTek). The chamber temperature was set to 37°C and the plate was shaken continuously in linear mode. At an OD600 of 0.3 Rhs2 expression was induced by the addition of 0.1% rhamnose and SciX expression was induced by the addition of 0.2% lactose. The OD600 of each well was read every 30 minutes for 16-18hrs. Experiments were performed in triplicate.

### Protein expression and purification of SciX

BL21(DE3)Gold cells containing SciX (pET21a+) were grown at 37°C in Lysogeny Broth (LB) media containing 100 μg/mL ampicillin until they achieved an absorbance at 600 nm of 0.6. Protein expression was induced by the addition of 1 mM Isopropyl β-D-1-thiogalactopyranoside (IPTG), and cells were further incubated for 20 h at 20°C. Next, the cells were centrifuged at 4100xg for 25 min and resuspended in wash buffer (50 mM Tris pH 8, 500 mM NaCl and 20 mM imidazole). The cells were then lysed using an Emulsiflex-C3 High Pressure Homogenizer (Avestin) and centrifuged at 35,000xg for 30 min. The supernatant was passed over a nickel-NTA agarose affinity chromatography gravity column (GoldBio) that had been equilibrated with the wash buffer. The protein was eluted with elution buffer (50 mM Tris pH 8, 500 mM NaCl and 500 mM imidazole). The N-terminal 6His-tag was cleaved using HRV-3C (human rhinovirus) protease at a dilution of 1:50, and the sample was dialyzed overnight in wash buffer at 4 °C. To remove undigested SciX, the sample was passed over a Ni-NTA agarose affinity chromatography gravity column (GoldBio) equilibrated in wash buffer. Subsequently, the dialyzed sample was concentrated to 2 ml and injected on a Superdex 75 Hi-Load (16/600) column (Cytiva) equilibrated in gel filtration buffer (50 mM sodium citrate pH 5.5, 250 mM NaCl and 1 mM beta-mercaptoethanol (β-ME) using an AKTA Pure (Cytiva). To confirm purity, all SciX samples were run on a 12% SDS-PAGE gel and visualized by staining with Coomassie dye.

### Nano-differential scanning fluorimetry

To discover an optimal buffer for SciX nano-differential scanning fluorimetry (nano-DSF) was used. SciX was expressed and purified by metal affinity chromatography as outlined in the protein expression and purification of SciX, but originally with a standard gel filtration buffer (50 mM Tris pH 7.5, 250 mM NaCl and 1 mM β-ME). Purified protein was diluted to 2 mg/ml using different buffers from the Solubility and Stability screens (Hampton Research). Initial hits were refined using buffers consisting of 50 mM Tris pH 8 (0, 50, 100, 150, 200 mM NaCl), 50 mM BisTris propane pH 8.5 (0, 50, 100, 150, 200 mM NaCl), 50 mM HEPES pH 7.5 (100, 150, 200 mM NaCl) and 50 mM sodium citrate pH 5.5, 250 mM NaCl. Protein samples were melted and Tms (melting temperatures) measured by increasing the temperature from 20 to 95 °C using a Prometheus NT.48 (NanoTemper).

### SEC-coupled multiple angle light scattering (SEC-MALS)

A Superdex 200 (10/300) increase column was equilibrated with gel filtration buffer containing 50 mM sodium citrate pH 5.5, 500 mM NaCl and 1 mM β-ME on an AKTA Pure (Cytiva). Purified SciX was concentrated to 10 mg/ml in the gel filtration buffer and spun in a 0.1 μm filter to remove aggregates before SEC-MALS analysis. SEC-MALS data was collected on a DAWN HELEOS II detector (Wyatt Technologies) coupled with an AKTA Pure with an in-line UV cell (Cytiva). SciX was injected after equilibration and detectors aligned and normalized with a 10 mg/ml BSA (bovine serum albumin) control (Sigma). All experiments were performed at 25°C. Analysis of the data was completed using ASTRA analysis software (Wyatt Technologies)

### Protein crystallization of SciX

Crystallization conditions for purified SciX were screened using commercially available screens (NeXtal, Molecular Dimensions), using a Crystal Gryphon robot (Art Robbins Instruments). Crystals were grown with crystallization conditions consisting of 40 mM potassium phosphate monobasic, 16% w/v PEG 8000 and 20% w/v glycerol at 20 mg/ml initially at 4°C. A 1:1 drop ratio using the sitting drop vapour diffusion method was used. Crystals were obtained after an incubation failure that heated the crystals to an unknown temperature before re-cooling to 4°C.

### X-ray Data collection and refinement

A SciX dataset was collected at the Advanced Light Source at Lawrence Berkeley National Laboratory using the ALS 8.3.1 Beamline. Data was integrated and scaled using XDS^77^ and merged with AIMLESS from the CCP4 program suit^78^. Phases were obtained using molecular replacement with Phaser^79^ using an AlphaFold model. The SciX structure was further refined using Coot^80^ and the Phenix package^81^, including TLS refinement^82^. Model statistics are listed in Table 1. Molecular visualizations and graphics were generated using ChimeraX^59^ and GraphPad Prism version 10.0.2.

### Isotopic labeling of SciX

Wildtype SciX was overexpressed in minimal media containing 20 mM sodium phosphate dibasic, 20 mM potassium phosphate monobasic, 10 mM sodium chloride, 1 mM magnesium sulphate, 0.1 mM calcium chloride, 10 mM iron chloride, 1 mg/ml of vitamin B1 and 1% glucose. Isotopic labeling was performed using 1 g/L of 15N ammonium chloride and 3 g/L 13C D-glucose. Additionally, micronutrients were added to the media including 3 μM ammonium molybdate tetrapeptide, 400 μM boric acid, 30 μM cobalt chloride, 10 μM copper sulphate, 80 μM manganese chloride and 10 μM zinc chloride. Cells were grown at 37 °C until they achieved an absorbance at 600 nm of 0.8. Protein expression was induced by 1 mM IPTG, and cells were further incubated for 20 h at 20°C.

### NMR spectroscopy

Triple resonance NMR spectra of SciX for backbone assignments were recorded at 25°C on a Bruker Avance II 700MHz spectrometer with a 0.5 mm TXI probe at the Regional Centre of NMR Spectroscopy at the University of Montreal, Quebec. Final samples of ^13^C and ^15^N labeled SciX consisted of 0.5 mM protein in 50 mM sodium citrate pH 5.5 and 250 mM NaCl with the addition of 5% D2O. Signals from the ^1^H, ^13^C, and ^15^N nuclei of SciX were detected and assigned by standard heteronuclear NMR experiments^83^. ^1^H-^15^N NOE relaxation data for SciX was recorded using a 500 MHz Bruker Avance III spectrometer and 5 mm BBFO probe at 25 °C following established methods^63^. NMR data was processed using NMRpipe^84^ and analyzed with Sparky (T. D. Goddard and D. G. Kneller, SPARKY 3, University of California, San Francisco)^85^. Backbone torsion angles were calculated from chemical shift assignments using TALOS^61,62^.

### Purification of Rhs2-EF-Tu

BL21(DE3)Gold cells containing Rhs2 D192A residues 137-246 (pET29b+) were grown at 37°C in Lysogeny Broth (LB) media containing 50 μg/mL kanamyacin until they achieved an absorbance at 600 nm of 0.6. Protein expression was induced by the addition of 1 mM Isopropyl β-D-1-thiogalactopyranoside (IPTG), and cells were further incubated for 20 h at 20°C. Next, the cells were centrifuged at 4100xg for 25 min and resuspended in wash buffer (50 mM Tris pH 8, 500 mM NaCl and 20 mM imidazole). The cells were then lysed using an Emulsiflex-C3 High Pressure Homogenizer (Avestin) and centrifuged at 35,000xg for 30 min. The supernatant was passed over a nickel-NTA agarose affinity chromatography gravity column (GoldBio) that had been equilibrated with the wash buffer. The protein was eluted with elution buffer (50 mM Tris pH 8, 500 mM NaCl and 500 mM imidazole). The elution was concentrated using an Amicon centrifugal concentrator with a 3 kDa cut-off and injected onto an SD75 Superdex Hi-Load (16/600) column (Cytiva). The size-exclusion buffer was 50 mM Tris pH 7.5, 250 mM NaCl and 1 mM β-ME. Fractions were visualized by Coomassie stained SDS-PAGE gel.

### Sample processing for mass-spectrometry

Bands containing the high molecular weight protein were cut from the Coomassie stained SDS-PAGE gel of purified Rhs2 D192A and stored in water at 4°C. Samples were delivered to the Manitoba Centre for Proteomics and Systems Biology core facility for processing and identification. Gel pieces were destained three times with 200 µL 100 mM NH₄HCO₃/acetyl nitrile (ACN) (1:1) for 10 min on a moderate vortex. Gel pieces were then dehydrated three times with 500 µL 100% ACN for 10 min until they shrank and turned opaque, and residual solvent was removed by SpeedVac. For protein reduction, gel pieces were rehydrated in 100 µL 10 mM dithiothreitol (DTT) in 25 mM NH₄HCO₃ and incubated at 57°C for 30 min. The solution was removed, and gel pieces were washed with 500 µL ACN for 10 min before residual solvent was removed by SpeedVac. For alkylation, gel pieces were rehydrated in 100 µL 55 mM IAA in 25 mM NH₄HCO₃ and incubated at room temperature in the dark for 30 min. The solution was removed, and gel pieces were washed with 500 µL ACN for 10 min before residual solvent was removed by SpeedVac. Gel pieces were rehydrated in 12.5 ng/µL trypsin in 50 mM NH₄HCO₃ at 4°C for 45 min. Digestion buffer was added as needed to cover the gel pieces, and digestion was carried out for 16 h at 37°C. Digests were collected by centrifuging the gel pieces at 800xg for 5 min. To increase peptide recovery, gel pieces were extracted once with water and three times with 5% formic acid in 50% ACN. All extracts were pooled, lyophilized, and resuspended in 0.5% formic acid. Ten percent of each sample was then loaded onto Evotip Pure EV2011 according to the manufacturer’s protocol for MS analysis.

### Liquid chromatography tandem mass spectrometry (LC/MS/MS)

An equivalent of 200 ng of peptides was analyzed by nanoflow LC-MS/MS using an Evosep One coupled to an Orbitrap Exploris 480 instrument (Thermo Fisher Scientific, Bremen, Germany) operated in positive mode using data-dependent acquisition. An Evosep EV1137 column was used to separate peptides (1.5 µm particles, 150 µm ID x 15 cm, Evosep) with 0.1% formic acid in water (mobile phase A) and 0.1% formic acid in acetonitrile (mobile phase B) using the preset 30 samples per day gradient and flow rate. Spray voltage was set to 2.15 kV, funnel RF level at 40%, and heated capillary at 275°C. The overall acquisition cycle of 1 second comprised one survey scan and MS/MS scans. Survey scans covering the mass range of 380–1500 m/z were acquired at a resolution of 60,000 (at m/z 200), with a normalized automatic gain control (AGC) target of 300% and a maximum ion injection time of 50ms. This was followed by MS2 acquisition at a resolution of 15,000, selected ions were fragmented at 30% normalized collision energy, with intensity threshold kept at 2e4. AGC target value for fragment spectra was set to standard with a maximum ion injection time set to auto and an isolation width set at 1.6 m/z. Dynamic exclusion of previously selected masses was enabled for 15 seconds, charge state filtering was limited to 2–5, peptide match was set to preferred, and isotope exclusion was enabled.

## DATA AVAILABILITY

The X-ray structure of *Salmonella* Typhimurium str. LT2 SciX has been deposited in the Protein Data Bank under accession code 9ZR2. The chemical shift assignments for SciX have been deposited in the Biological Magnetic Resonance Bank under accession code 53755.

## SUPPORTING INFORMATION

This article contains supporting information.

## AUTHOR CONTRIBUTIONS

N.L.C. and G.P. conceived the study and contributed to the experimental design. N.L.C. expressed, purified, and crystallized proteins and performed all bacterial assays. S.G. performed molecular cloning of point variants. D.D. configured and optimized the collected NMR relaxation data. N.L.C. solved the crystal structure and N.L.C. and G.P. processed and analyzed all NMR data. N.L.C. and G.P. performed structural modeling and analysis. N.L.C. and G.P. wrote the paper. All authors provided feedback on the manuscript.

## ACKNOWLEDGEMENTS

We would like to thank the laboratory of Jörg Stetefeld at the University of Manitoba for SEC-MALS instrument access and Markus Meier for assistance with data collection and processing. We also thank Brian L. Mark at the University of Manitoba and Beamline 8.3.1 at the Advanced Light Source for assistance with X-ray data collection. We thank Regional Centre of NMR spectroscopy at the University of Montreal for NMR access and Normand Cyr for helping to collect and process NMR spectra. We thank both Ying Lao and Rene Zahedi at the Manitoba Centre for Proteomics and Systems Biology for help with mass-spectrometry data collection and analysis.

## FUNDING AND ADDITIONAL INFORMATION

This work was supported by a Natural Sciences and Engineering Research Council of Canada (NSERC) Discovery Grant RGPIN-2025-05624 to G.P., a Canadian Foundation for Innovation (CFI) grant 37841 to G.P, and a University of Manitoba Research Grants Program (URGP) grant to G.P. N.L.C. was supported by a University of Manitoba Graduate Fellowship and S.G. was supported by a Research Manitoba Graduate Studentship and a University of Manitoba Graduate Fellowship.

