## Supplemental_data for "Molecular characterization of the effector-immunity pair Rhs2-SciX from *Salmonella* Typhimurium"

Running title: Rhs2 and SciX are homologous to BECR family effector-immunity pairs

**Key words:**
Type VI secretion system, effector, immunity protein, X-ray crystallography, NMR spectroscopy, *Salmonella* Typhimurium

*To whom correspondence should be addressed: G.P.


List of supplementary information
Table S1
Figures S1-S7

**Table S1: Data table for SEC-MALS of SciX**

| | |
|---|---|
| **Hydrodynamic radius (Q) moments (nm)** | |
| **rh (Q) (avg)** | 1.435 |
| **General (mL/(mg cm))** | |
| **UV Ext Coef. (mL/(mg cm))** | 1.834 |
| **Masses** | |
| **Injected Mass (µg)** | 1464.00 |
| **Calculated Mass (µg)** | 712.72 |
| **Mass Recovery (%)** | 48.7 |
| **Mass Fraction (%)** | 100.0 |
| **Molar mass moments (g/mol)** | |
| **Mn** | 16840 |
| **Mp** | 18790 |
| **Mv** | N/A |
| **Mw** | 16950 |
| **Mz** | 17050 |
| **Mz+1** | 17160 |
| **Polydispersity** | |
| **MW/Mn** | 1.006 |
| **Mz/Mn** | 1.013 |
| **Light scattering peak statistics** | |
| **Retention Time (mL)** | 17.922 |
| **Mean (mL)** | 17.884 |
| **Refractive index peak statistics** | |
| **Retention Time (mL)** | 17.954 |
| **Mean (mL)** | 17.873 |
| **UV peak statistics** | |
| **Retention Time (channel 1) (mL)** | 17.957 |
| **Mean (channel 1) (mL)** | 17.868 |

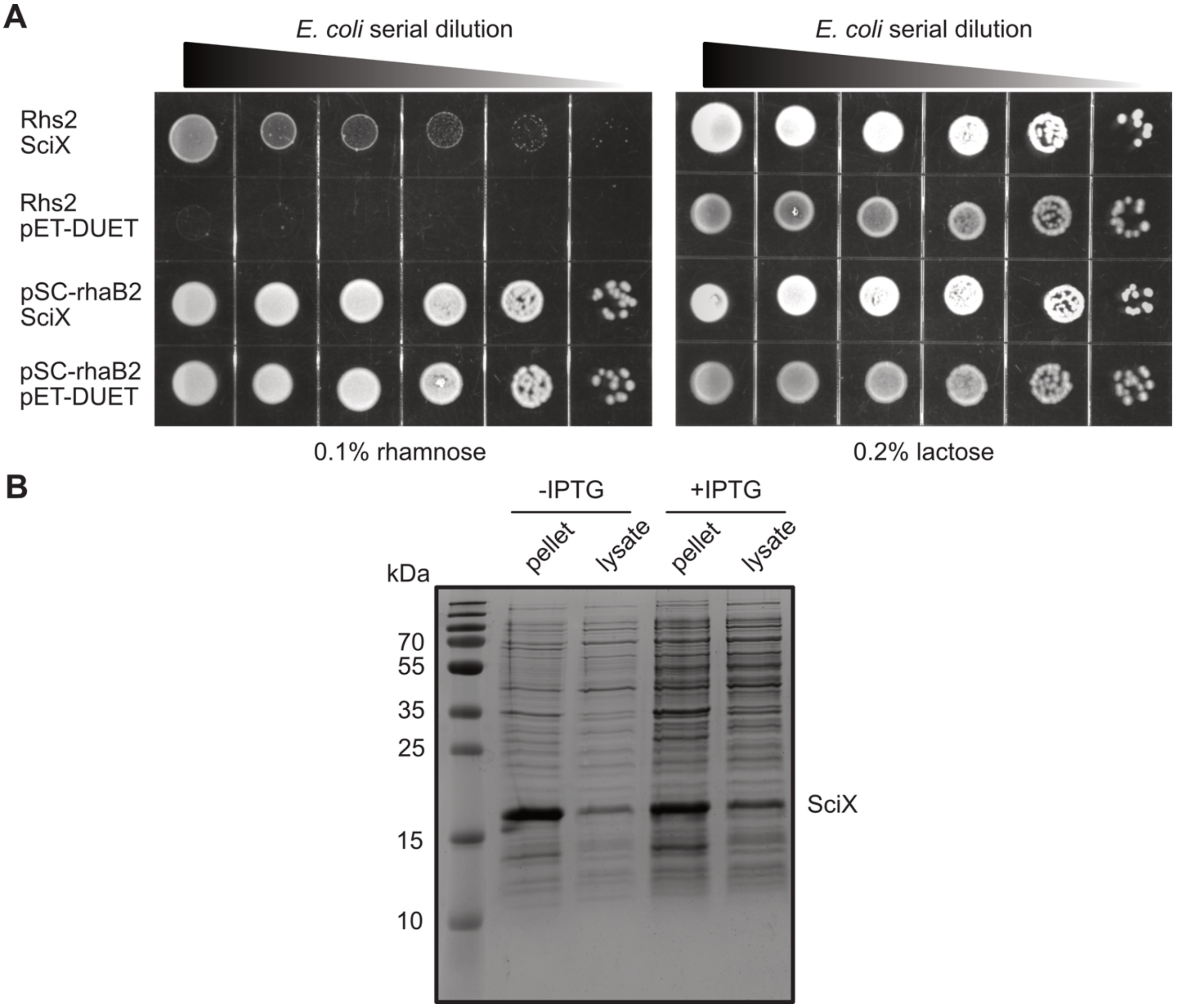


**Figure S1: Bacterial toxicity assay controls for Rhs2-SciX A)** Rhs2-SciX effector-immunity toxicity assays by end-point plate. Serial dilutions of *E. coli* co-transformed with *rhs2* pSC-rhaB2 (Rhs2) and *sciX* pETDUET-1 (SciX) or with the empty vectors pSC-rhaB2 and pETDUET-1. Left: induction with 0.1% rhamnose for Rhs2 expression, Right: induction with 0.2% lactose expression for SciX. **B)** Induction trail of SciX under the control of a T7 promoter. SciX shows some protein expression even in the absence of the inducer ITPG. SciX is labeled by approximate molecular weight of 17 kDa.

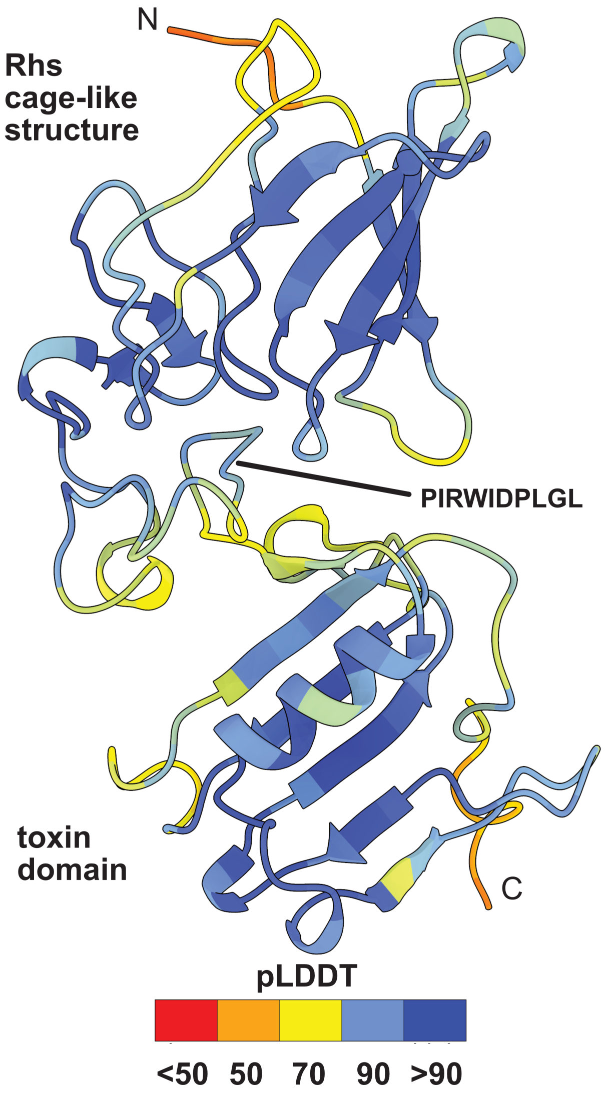


**Figure S2: AlphaFold model of Rhs2.** AlphaFold3 model of Rhs2 colored by pLDDT score. Red is low confidence and dark blue is high confidence. Overall pTM is 0.68.The Rhs-cage structure and toxin domain are labeled in addition to the conserved T6SS Rhs-toxin cleavage motif between domains.

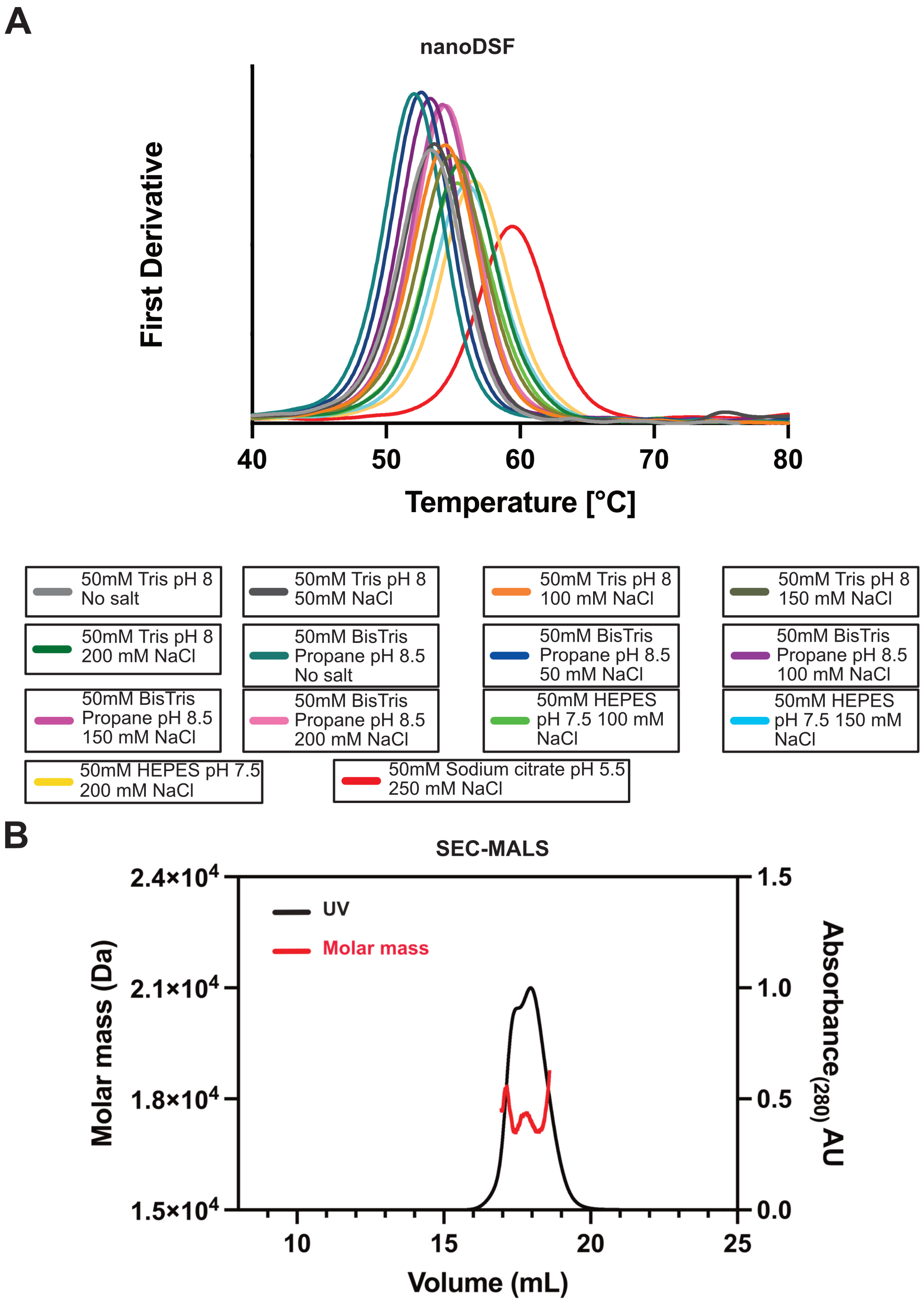


**Figure S3: Biophysical analysis of SciX. A)** SciX buffer optimization by nano-differential scanning fluorimetry. The peak is the first derivative of the ratio of tryptophan fluorescence (330/350) and is the melting temperature of SciX in each buffer. All buffers are listed and plotted by color. **B)** SEC-MALS trace of SciX showing UV and calculated molar mass. Data was analyzed using ASTRA software.

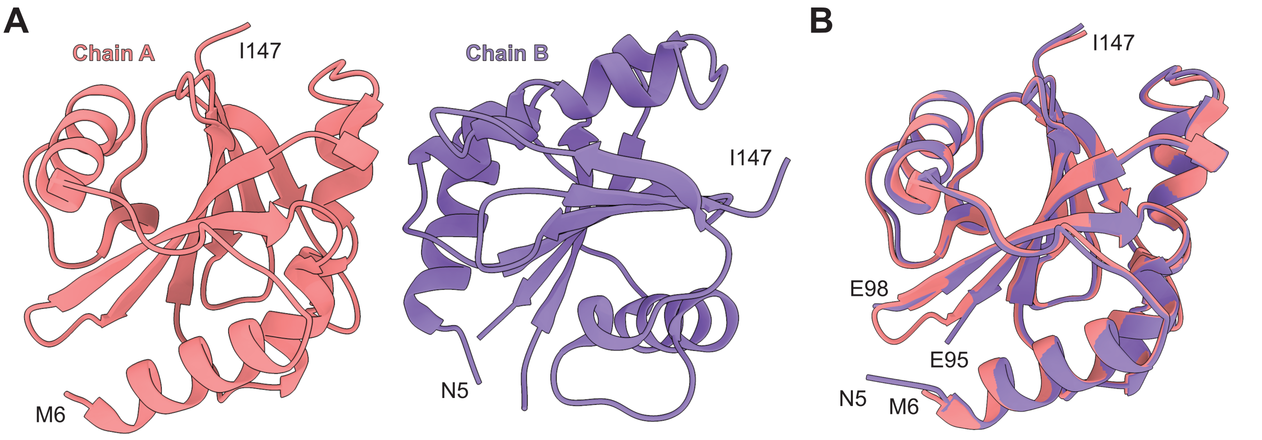


**Figure S4: Asymmetric unit contents of SciX crystal structure. A)** Chain A is shown in pink and Chain B in purple. **B)** Structural alignment of both chains has a rmsd of 0.5 Å indicating identical conformations.

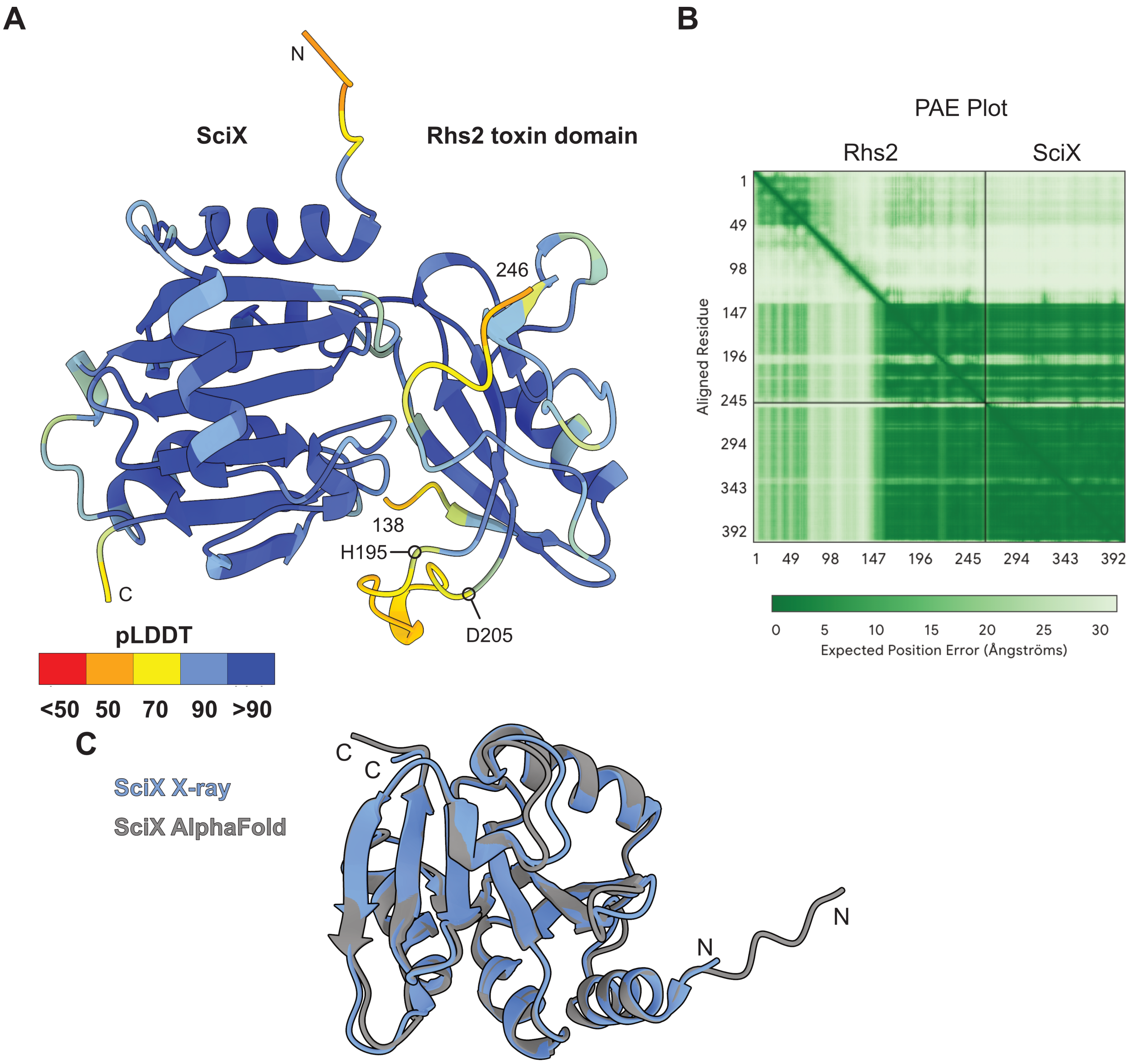


**Figure S5: AlphaFold model of an Rhs2-SciX complex. A)** AlphaFold3 model of an Rhs2-SciX complex colored by pLDDT score. Only the Rhs2 toxin domain bound to SciX is shown. Red is low confidence and dark blue is high confidence. **B)** PAE plot of the predicted Rhs2-SciX complex. Rhs2 and SciX are labeled. **C)** Overlay of the SciX AlphaFold3 model (grey) and the SciX crystal structure (blue).

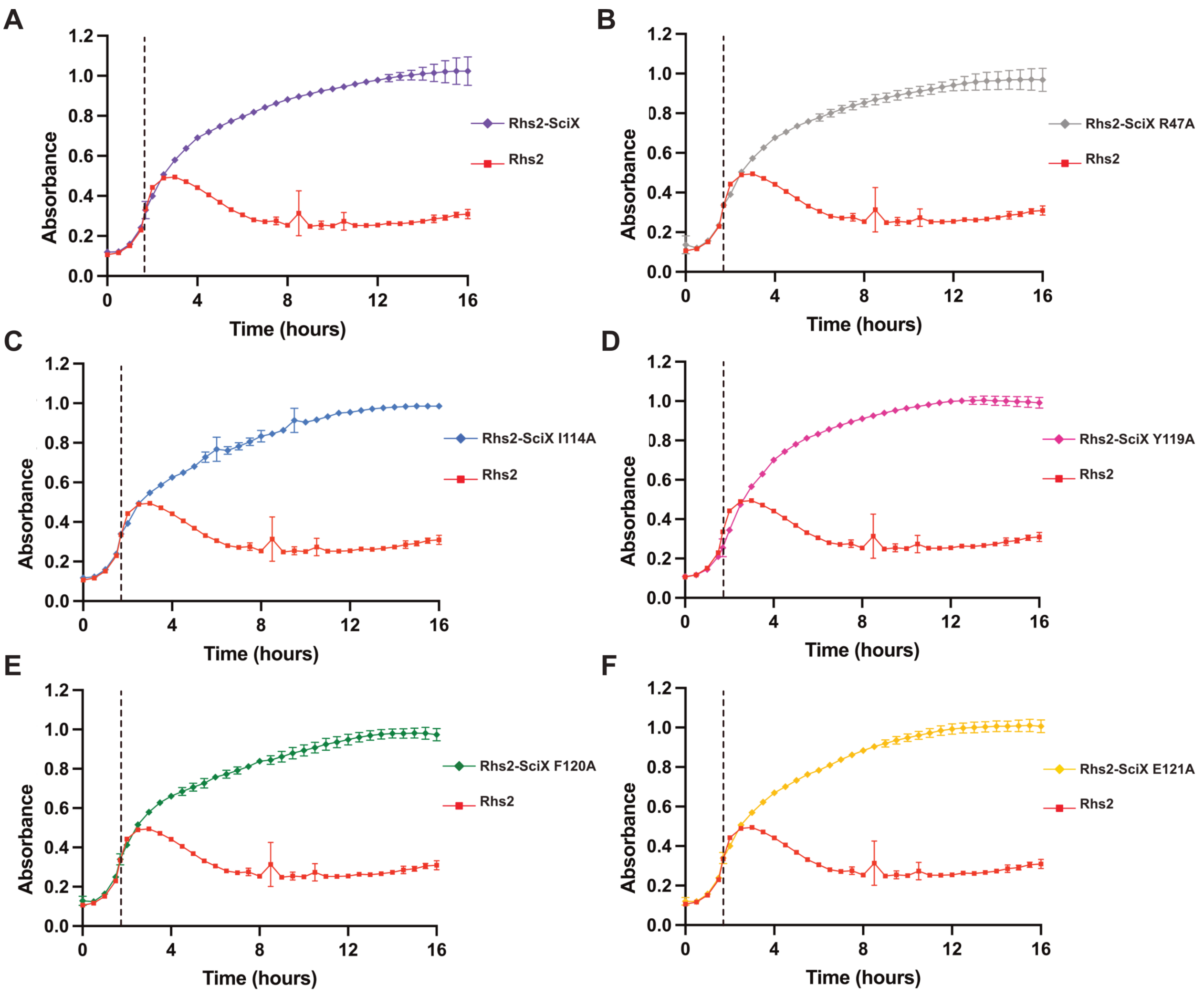


**Figure S6: Bacterial growth assays of Rhs2 and SciX variants. A)** Wild-type SciX and Rhs2 redrawn from previous figures. B) Rhs2 and SciX R47A C) Rhs2 and SciX I114A D) Rhs2 and SciX Y119A E) Rhs2 and SciX F120A F) Rhs2 and SciX E121A. Samples were induced with 0.1% rhamnose and 0.2% lactose at an of OD600 = 0.3. Point of induction is indicated by dotted line. The Rhs2 curve (red) is repeated from previous figures for comparison. Samples were run in triplicate.

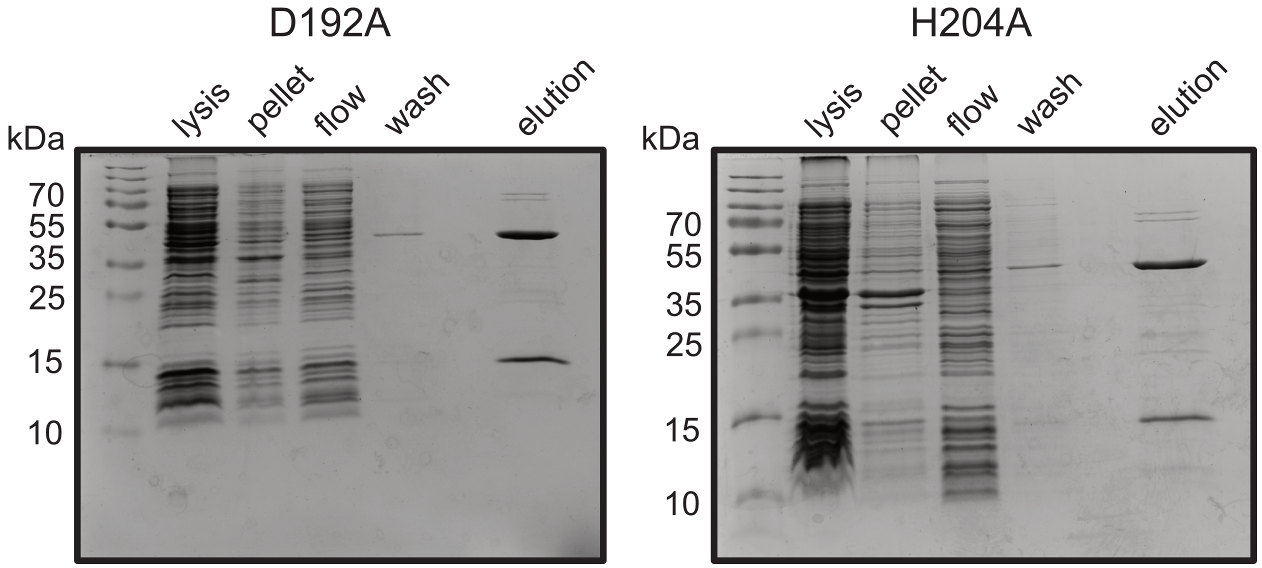


**Figure S7: Affinity-tag purification of Rhs2 toxin domain variants D192A and H204A.** Purification stages of each variant are shown by Coomassie stained SDS-PAGE gel. Each lane is labeled by purification step. The final elution band shows the Rhs2 toxin domain and a larger 45 kDa protein.
